# Kinome screening identifies integrated stress response kinase EIF2AK1 / HRI as a negative regulator of PINK1 mitophagy signaling

**DOI:** 10.1101/2023.03.20.533516

**Authors:** Pawan K. Singh, Shalini Agarwal, Ilaria Volpi, Léa P. Wilhelm, Giada Becchi, Andrew Keenlyside, Thomas Macartney, Rachel Toth, Adrien Rousseau, Glenn R. Masson, Ian G. Ganley, Miratul M. K. Muqit

## Abstract

Loss-of-function mutations of the PINK1 kinase cause familial early-onset Parkinson’s disease (PD). PINK1 is activated upon mitochondrial damage to phosphorylate Ubiquitin and Parkin to trigger removal of damaged mitochondria by autophagy (mitophagy). PINK1 also indirectly phosphorylates a subset of Rab GTPases including Rab8A. We have performed an siRNA screen targeting all human Ser/Thr kinases in HeLa cells and discovered that knockdown of the eukaryotic translation initiation factor 2-alpha kinase 1 (EIF2AK1), also known as heme-regulated inhibitor (HRI) kinase, a branch of the integrated stress response (ISR), selectively enhances mitochondrial depolarization-induced stabilization of PINK1 and increased phosphorylation of ubiquitin and Rab8A. We confirm our findings in multiple human cell lines, including SK-OV-3, U2OS and ARPE-19 cells. Knockdown of the upstream mitochondrial-cytosol relay component, DELE1, enhanced PINK1 stabilisation and activation similar to EIF2AK1 knockdown. Strikingly, we demonstrate that the small molecule ISR inhibitor, ISRIB, also enhances PINK1 activation and signaling under conditions of mitochondrial damage. Using the *mito*-QC mitophagy reporter in human cells, we observe that EIF2AK1 knockdown or ISRIB treatment significantly enhances PINK1-dependent mitophagy but does not alter deferiprone-induced mitophagy. Our findings indicate that the DELE1-EIF2AK1 ISR signaling relay is a negative regulator of PINK1-dependent mitophagy and suggest that inhibitors of DELE1-EIF2AK1 and/or ISRIB analogues could have therapeutic benefits in PD and related disorders.

## INTRODUCTION

Autosomal recessive mutations of genes encoding the mitochondrial protein kinase PTEN-induced kinase 1 (PINK1) and RING-IBR-RING (RBR) Ubiquitin E3 ligase Parkin are causal for familial early-onset Parkinson’s disease (PD) [1, 2]. Upon mitochondrial depolarisation-dependent damage that can be induced in cells by chemical uncouplers (e.g. Oligomycin/Antimycin A (OA)), PINK1 undergoes stepwise activation with (1) protein stabilisation (2) recruitment to the Translocase of Outer Membrane (TOM) complex at the outer mitochondrial membrane (OMM); (3) dimerization; (4) trans-autophosphorylation of residue Serine228 (S228) and (5) stabilisation of loop insertion 3 within its catalytic domain that enables the recognition of substrates Ubiquitin and Parkin [3–8]. PINK1 directly phosphorylates an equivalent Serine 65 (S65) residue found in Ubiquitin and Parkin resulting in maximal activation of Parkin via a feed-forward mechanism triggering Ubiquitin-dependent removal of damaged mitochondria by autophagy (mitophagy) [9–11]. We have also found that PINK1 activation leads to the phosphorylation of a subset of Rab GTPases including Rab8A at a highly conserved Serine111 residue located within its RabSF3 motif [12, 13]. The mechanism of Rab protein phosphorylation by PINK1 is indirect, suggesting the role of an unknown intermediate kinase [12, 13]. Furthermore, Rab8A Ser111 phosphorylation (Rab8A pS111) is a robust biomarker of PINK1 activation in cells and is abolished in human PINK1 knockout cells or cells expressing PD-associated mutants of PINK1 [4, 12].

The integrated stress response (ISR) pathway consists of four branches: EIF2AK1/HRI (heme-regulated inhibitor), EIF2AK2/PKR (protein kinase R), EIF2AK3/PERK (protein kinase R-like ER kinase) and EIF2AK4/GCN2 (general control non-derepressible 2), each responding to distinct stress signals [14–17]. Upon sensing stress, ISR kinases phosphorylate eukaryotic translation initiation factor 2α (eIF2α) resulting in the downregulation of protein synthesis and induction of the transcription factor ATF4 (activating transcription factor 4) [14]. Recent pioneering work from the Jae and Kampmann labs has identified that upon mitochondrial stress, the mitochondrial localised protein DAP3-binding cell death enhancer 1 (DELE1) is cleaved by the mitochondrial protease OMA1, and the C-terminal cleaved fragment of DELE1 is released into the cytosol, where it binds and activates EIF2AK1, leading to downstream ISR signaling [18, 19]. Previous studies have linked ISR with PD pathology [20] although no studies have directly assessed EIF2AK1 or DELE1 protein levels in PD tissues. Furthermore, no genetic variants in EIF2AK1 or DELE1 have been found as a risk factor for PD, although genetic studies have reported that EIF2AK3/PERK gene variants are a risk factor for the related neurodegenerative disorder, Progressive Supranuclear Palsy [21].

An open question in the field is whether PINK1 is controlled by other kinase signaling pathways and networks. Herein, we have undertaken a genetic siRNA screen targeting all known human Ser/Thr kinases using endogenous PINK1-phosphorylated Rab8A at Ser111 as a readout in HeLa cells and found that siRNA-mediated knockdown of the integrated stress response kinase, EIF2AK1 (that encodes Heme Regulated Inhibitor; HRI) leads to a significant increase in Rab8A pS111 levels. We demonstrate that EIF2AK1 knockdown leads to increased stabilization and activation of PINK1 as measured by ubiquitin Ser65 phosphorylation (Ub pS65) following mitochondrial depolarisation. We further show that siRNA mediated knockdown of DELE1 can also increase PINK1 stabilisation and activation consistent with its role in activating EIF2AK1 under mitochondrial stress. These results are corroborated by the striking observation that the small molecule ISR inhibitor ISRIB can also promote PINK1 stabilisation and activation under conditions of mitochondrial depolarisation similar to EIF2AK1 knockdown. Using the *mito*-QC mitophagy reporter [22, 23], we also show that EIF2AK1 knockdown or ISRIB treatment enhances PINK1-dependent mitophagy but does not alter deferiprone (DFP)-induced mitophagy. Our results suggest that inhibition of the ISR by targeting DELE1-EIF2AK1 or by ISRIB and related analogues offer novel therapeutic strategies against PD.

## RESULTS

### Genetic siRNA screen for identification of kinases that regulate endogenous PINK1-dependent Rab8A phosphorylation

To identify protein kinases that regulate endogenous PINK1-dependent Rab8A phosphorylation, we performed a siRNA screen targeting all Ser/Thr kinases in HeLa cells. We assembled and transfected a library of 428 siRNA pools (Horizon Dharmacon) targeting each kinase component for 72 hr with mitochondrial depolarisation by adding 1μM Oligomycin/10μM Antimycin A (OA) in the last 20 h of knockdown (to stabilize and activate PINK1) before cell lysis (Fig. 1A). To standardize the screen, we also included untransfected cells (UT), mock transfected (no siRNA) and cells transfected with non-targeting (NT) siRNA as negative control and PINK1 siRNA as a positive control. As a readout of PINK1 kinase activity we monitored Ser111-phosphorylation of Rab8A deploying a previously developed phospho-specific antibody that detects endogenous Rab8A pS111 by immunoblot analysis [11]. Rab8A pS111 and total Rab8A levels were quantified in parallel employing a multiplex immunoblot assay and the ratio of Rab8A pS111 *versus* total Rab8A was used to calculate the degree of Rab8A phosphorylation in each cell lysate (Fig. 1A). Levels of total PINK1, GAPDH and OPA1 (cleaved OPA1 is a readout of mitochondrial depolarization) were also analyzed. In addition, immunoblotting of select kinase components of the siRNA library was performed to assess overall efficiency including BCR, TRIM28, RPS6KB2, BRD2, MAP3K7, CDC7, ROCK1, EEF2K, CAMK1D, GRK2, CSNK1A1, CSNK2A1, MAPK3, EIF2AK4, PBK/SPK, MAP3K5, MAP2K2, MAP2K6 and MAP2K7 (fig. S1-S4). Strikingly, knockdown of EIF2AK1 enhanced Rab8A pS111 over 1.8-fold (Fig. 1B). This was associated with an over 2-fold increase in total PINK1 levels (Fig. 1C). Hits that enhanced PINK1 levels greater than 1.5-fold included SIK1 and CDK13; however, these did not significantly alter levels of Rab8A pS111. Conversely, we did not detect any hits that significantly reduced phospho-Ser111 Rab8A similar to PINK1 (Fig. 1B). Knockdown of a number of kinases mildly reduced Rab8A pS111 but were not significant (Table S1).

**Fig. 1.**
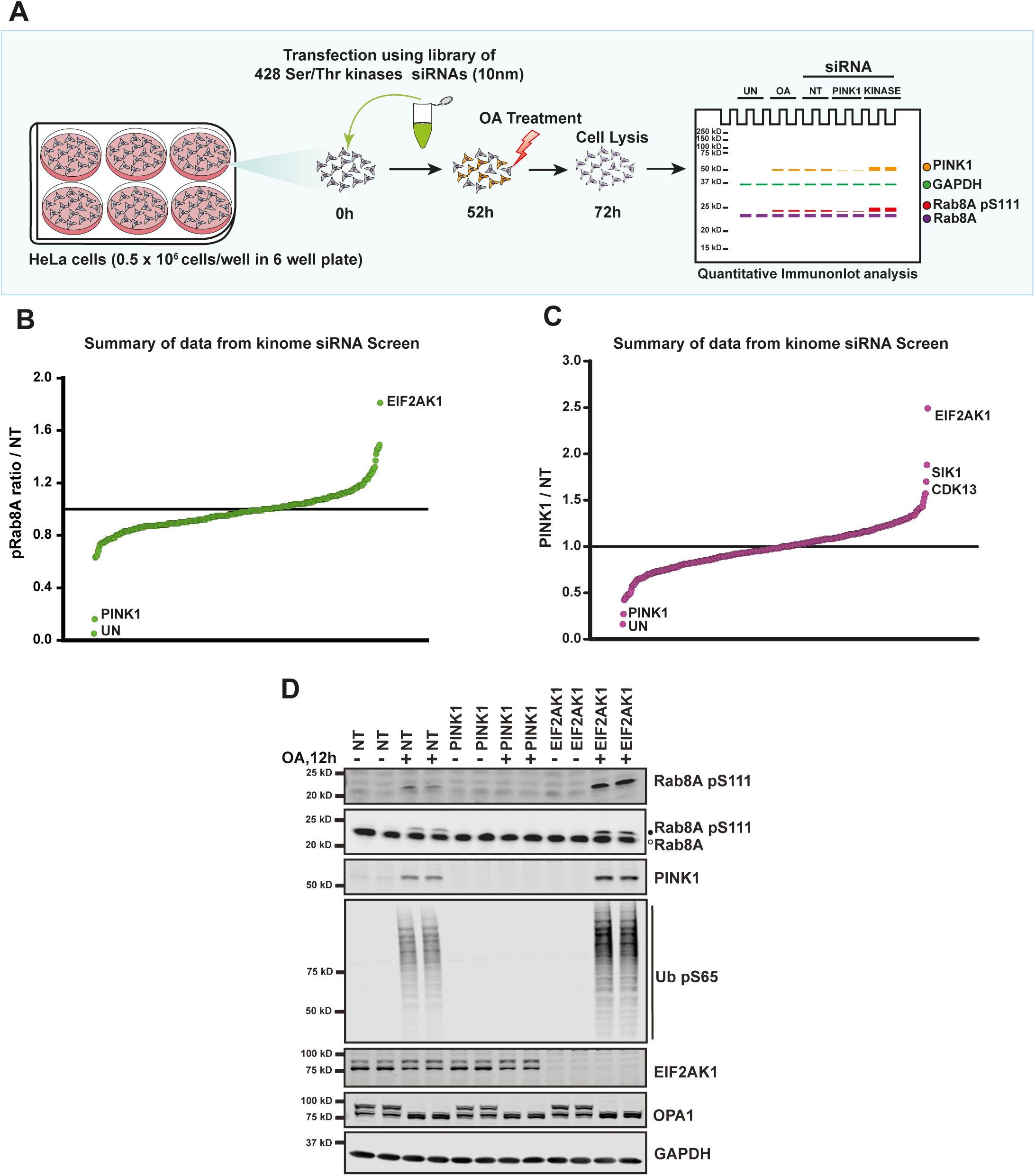
siRNA screen to identify kinases that regulate endogenous PINK1-dependent Rab8A phosphorylation. (**A**) Schematic of the siRNA knockdown screens employed in this study. HeLa cells seeded in 6-well plates at 0.5*10^6^ cells/well were transfected with siRNA pools (Dharmacon) for 72 hr targeting 428 Ser/Thr Kinases with Oligomycin (1 μM)/Antimycin A (10 μM) (OA) treatment for the last 20 hr of siRNA targeting. Cells were lysed and immunoblotted for indicated antibodies and developed using the LI-COR Odyssey CLx Western Blot imaging system for quantitative analysis. (**B**) Summary of data from the kinome screen. The calculated ratio of Rab8A pS111/total Rab8A relative to non-targeting (NT) siRNA control, ranked from the highest increase in Rab8A phosphorylation to the strongest decrease (mean of the two replicates, SD values not shown on the chart, calculated using the Licor Image Studio software, screening blots provided in Supplementary file 1-4). (**C**) As in (**B**), the calculated ratio of PINK1/total GAPDH relative to NT siRNA control, ranked from highest increase in PINK1 levels to the strongest decrease. (**D**) HeLa cells were transfected with siRNA for 72 hr with the top hit from the screen, EIF2AK1 along with the siRNA for PINK1 and NT siRNA control and OA treatment was done for the last 12 hr of siRNA targeting. Cells were then lysed and immunoblotted with Rab8A pS111, Rab8A, PINK1, Ub pS65, EIF2AK1, OPA1 and GAPDH antibodies and analyzed as described above.

To confirm that siRNA-mediated knockdown of EIF2AK1 enhances PINK1-mediated Rab8A pS111 levels, we generated a sheep polyclonal antibody against total human EIF2AK1 using recombinant full-length GST-EIF2AK1 protein expressed and purified from *E. coli* (fig. S5A). This antibody specifically recognizes EIF2AK1, but not the related EIF2AK4 by immunoblotting (fig. S5B and C). Using this antibody, we were able to assess the degree of EIF2AK1 knockdown associated with PINK1 stabilisation/activation under our siRNA conditions alongside NT and PINK1 siRNA (Fig. 1D). Notably we did not observe any effect of EIF2AK1 siRNA on PINK1 levels in basal/unstimulated cells (Fig. 1D). Consistent with the screen results, upon mitochondrial depolarization induced by OA, we observed in EIF2AK1 knockdown cells a robust increase in PINK1 protein stabilization and Rab8A Ser111 phosphorylation (Fig. 1D). In addition, we observed that running lysates on 12% Tris-Glycine gels enabled visualization of an electrophoretic mobility band-shift of Rab8A that accompanied Rab8A pS111 and this was abolished by PINK1 siRNA knockdown and enhanced by EIF2AK1 knockdown, providing a facile readout to monitor PINK1-dependent Rab8A pS111 (Fig. 1D). To confirm that the increased PINK1 stabilisation is associated with increased PINK1 activation, we also measured levels of PINK1-dependent Ub pS65 and similarly found that EIF2AK1 siRNA knockdown increased levels of Ub pS65 (Fig. 1D). To verify the specificity of siRNA towards EIF2AK1, we performed similar experiments with independent pooled siRNA (Sigma) as well as single siRNAs (Dharmacon) for EIF2AK1 and observed that all led to enhanced PINK1 stabilisation, and PINK1-phosphorylated Rab8A as determined by Rab8A bandshift (fig. S6).

### Genetic validation studies confirm that specific knockdown of the ISR kinase, EIF2AK1, enhances PINK1 stabilization and activation in response to mitochondrial damage

We next compared the effect of EIF2AK1 siRNA knockdown with the knockdown of the other three eIF2α kinases, EIF2AK2, EIF2AK3, and EIF2AK4. Our observations revealed that only EIF2AK1 knockdown selectively enhanced PINK1 stabilisation and activation as judged by increased levels of Ub pS65 and Rab8A pS111 (band-shift) (Fig. 2A-D). Furthermore, we also observed a robust induction of ATF4 with OA treatment that was abolished by EIF2AK1 knockdown but not following EIF2AK2/3/4 knockdown consistent with previous studies (Fig. 2A) [18, 19]. To determine whether regulation of PINK1 stabilisation by EIF2AK1 is generalizable to other cells, we screened a panel of human cell lines including SK-OV-3 ovarian cancer cells that express high levels of endogenous PINK1 and Parkin [4, 24]; U2OS cells that express moderate PINK1 and low levels of Parkin; and ARPE-19 retinal pigment epithelial cells that have moderate PINK1 expression and no Parkin similar to HeLa cells (fig. S7A-C). Upon treatment of OA, we observed in all 3 cell lines, that siRNA-mediated knockdown of EIF2AK1 enhanced PINK1 stabilisation and activation as measured by Ub pS65 and Rab8A pS111 and this was also associated with inhibition of OA-induced ATF4 expression (fig. S7A-C).

**Fig. 2.**
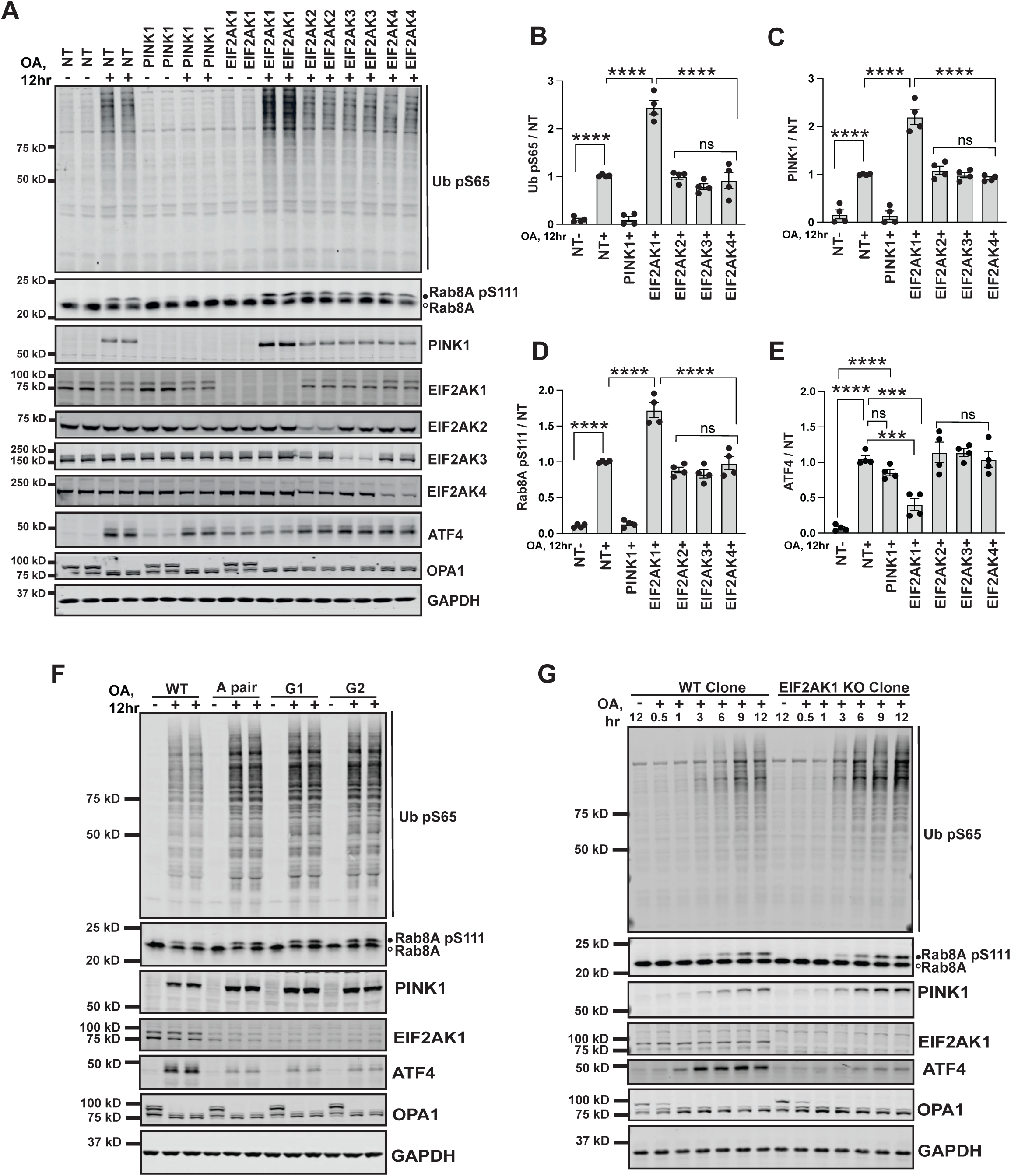
siRNA knockdown and CRISPR knockout of the full-length EIF2AK1 in HeLa cells enhances PINK1 stabilisation and activation. (**A**) HeLa cells either treated with non-targeting control siRNA pool (NT), siRNA pool targeting PINK1, EIF2AK1, EIFF2AK2, EIF2AK3, EIF2AK4, for 72 hr were treated with OA in the last 12 hr. Cells were lysed and immunoblotted for, Ub pS65, RAB8A, PINK1, EIF2AK1, EIFF2AK2, EIF2AK3, EIF2AK4, OPA1, ATF4, GAPDH and the immunoblots were developed using the LI-COR Odyssey CLx Western Blot imaging system. Quantification shown on right as (**B**) Quantification of Ub pS65/GAPDH ratio normalised to the ratio in NT (+OA) samples using the Image Studio software. (**C**) Quantification of PINK1/GAPDH ratio normalised to the ratio in NT (+OA) samples. (**D**) Quantification of Rab8A pS111/GAPDH ratio normalised to the ratio in NT (+OA) samples. (**E**) Quantification of ATF4/GAPDH ratio normalised to the ratio in NT (+OA) samples. (**F**) Wild-type HeLa cells and HeLa cells transfected with three different sets of CRISPR/Cas9 EIF2AK1 guide RNAs (A pair, G1 and G2) were treated with OA for 12 hr. Cells were lysed and immunoblotted for Ub pS65, PINK1, Rab8A, EIF2AK1, ATF4, OPA1, GAPDH and developed using the LI-COR Odyssey CLx Western Blot imaging system. (Quantification shown as fig. S8 A-C). (**G**) Selected EIF2AK1 knockout clone A3 and wild-type HeLa cells treated with OA for 0.5, 1, 3, 6, 9 and 12 hr as above were analyzed for Ub pS65, PINK1, Rab8A, EIF2AK1, ATF4, OPA1 and GAPDH. Data information: (**B**,**C**,**D** and **E**) All data are mean ± SEM; Statistical significance is displayed as *P ≤ 0.05; **P ≤ 0.01; ***P ≤ 0.001; ****P ≤ 0.0001; ns, not significant. n = 4 technical replicates (2 biological replicates), one-way ANOVA, Tukey’s multiple comparisons test.

EIF2AK1/HRI was initially implicated in a specialized role in erythrocyte (red blood cell) development whereby it is activated upon falling heme concentrations via its N-terminal heme-binding domain [25, 26]. However, it is now established that EIF2AK1 is widely expressed and responds to multiple types of signals including mitochondrial dysfunction, oxidative stress, and heat shock [15, 18, 19]. Mitochondrial dysfunction due to protein misfolding or mitochondrial depolarization induces up-regulation of the master transcriptional factor, ATF4 and expression of ATF4-target genes [27–29]. Consistent with this we observed a robust induction of ATF4 in OA-treated cells, which was abolished by EIF2AK1 knockdown (Fig. 2A, E and fig. S7A-C).

To further validate the role of EIF2AK1, we employed CRISPR/Cas9 gene-editing to knock out EIF2AK1 in HeLa cells. CRISPR guides (single guide1, guide 2, and guide pair A) were transfected into HeLa cells and immunoblot analysis of the CRISPR EIF2AK1 knockdown cellular pool lysates for all the guides demonstrated increased PINK1 stabilisation and activation as measured by Ub pS65 and Rab8A pS111 (Fig. 2F and fig. S8A-C). Following single-cell sorting and screening, we isolated 2 independent clones from guide pair A that were confirmed by sequencing and immunoblotting to be homozygous for loss-of-function mutations in the EIF2AK1 gene (fig. S9A-E). EIF2AK1 knockout clones, A2 and A3, were next treated with DMSO or OA for 12 hr to induce mitochondrial depolarization. Immunoblot analysis of whole cell lysates under basal conditions did not demonstrate any change in PINK1 levels or activity in EIF2AK1 knockout cells compared to controls consistent with the siRNA studies (fig. S9F). However, upon OA treatment, there was a robust increase in PINK1 stabilisation and activation as judged by Ub pS65 in both EIF2AK1 knockout clones compared to the wild-type control cell (fig. S9F). We next determined the time course of PINK1 stabilisation, Rab8A pS111 (band-shift) and Ub pS65 in the EIF2AK1 clone A3 knockout cells following OA treatment. We observed PINK1 protein levels becoming stable after 3 hr of OA treatment, in the wild-type control cells, associated with the accumulation of Ub pS65 and Rab8A pS111 and all three readouts were enhanced in EIF2AK1 knockout cells thereby validating the siRNA studies (Fig. 2G). We also observed a clear time-dependent up-regulation of ATF4 following OA treatment from 1 to 12 hrs (Fig. 2G). This up-regulation was abolished in EIF2AK1 knockout cells, indicating that EIF2AK1 is the primary ISR kinase activated by mitochondrial dysfunction in our cell system (Fig. 2G).

We also noted that total EIF2AK1 levels were variably reduced in OA stimulated cells as compared to basal/untreated cells (e.g. Fig. 2A). To address whether this effect is mediated by the activation of PINK1, we compared EIF2AK1 protein levels in wild-type and PINK1 knockout S-HeLa cells across a timecourse of OA stimulation. We observed a mild decrease in EIF2AK1 protein levels following OA treatment and there was no significant difference between the wild-type and PINK1 knockout cells (fig. S10). Therefore, OA-induced lowering of EIF2AK1 is PINK1 independent and recently it was reported that silencing factor of the integrated stress response (SIFI), an E3 ligase complex, mutated in early onset ataxia and dementia can regulate the degradation of EIF2AK1, to promote survival of cells undergoing mitochondrial import stress [30].

### DELE1-EIF2AK1 signaling relay negatively regulates PINK1

Previous studies had revealed that upon mitochondrial damage-induced activation of the mitochondrial protease OMA1, an inner mitochondrial membrane protein DELE1 is cleaved by OMA1 leading to accumulation of a shortened C-terminal fragment of DELE1 in the cytosol [18, 19]. This cleaved DELE1 fragment then oligomerizes in the cytoplasm and directly binds to and activates EIF2AK1, thereby initiating the ISR pathway [18, 19]. When exposed to oligomycin-induced mitochondrial stress, the cleaved DELE1 fragments are preferentially tagged by the SIFI complex and degraded in a manner similar to EIF2AK1 [30]. To investigate the role of DELE1 in PINK1 activation and signaling, we performed siRNA mediated knockdown of DELE1 alongside EIF2AK1 in HeLa cells (Fig. 3). We observed that following OA treatment, DELE1 knockdown led to a similar enhancement of PINK1 stabilisation and activation as EIF2AK1 knockdown (Fig. 3A-D). We also observed a substantial reduction in ATF4 levels in DELE1 knockdown cells, akin to the reduction seen in EIF2AK1 knockdown cells (Fig. 3A, E). These findings highlight a major role of the DELE1-EIF2AK1-ISR signaling relay in negatively regulating mitochondrial damage-induced activation of PINK1.

**Fig. 3.**
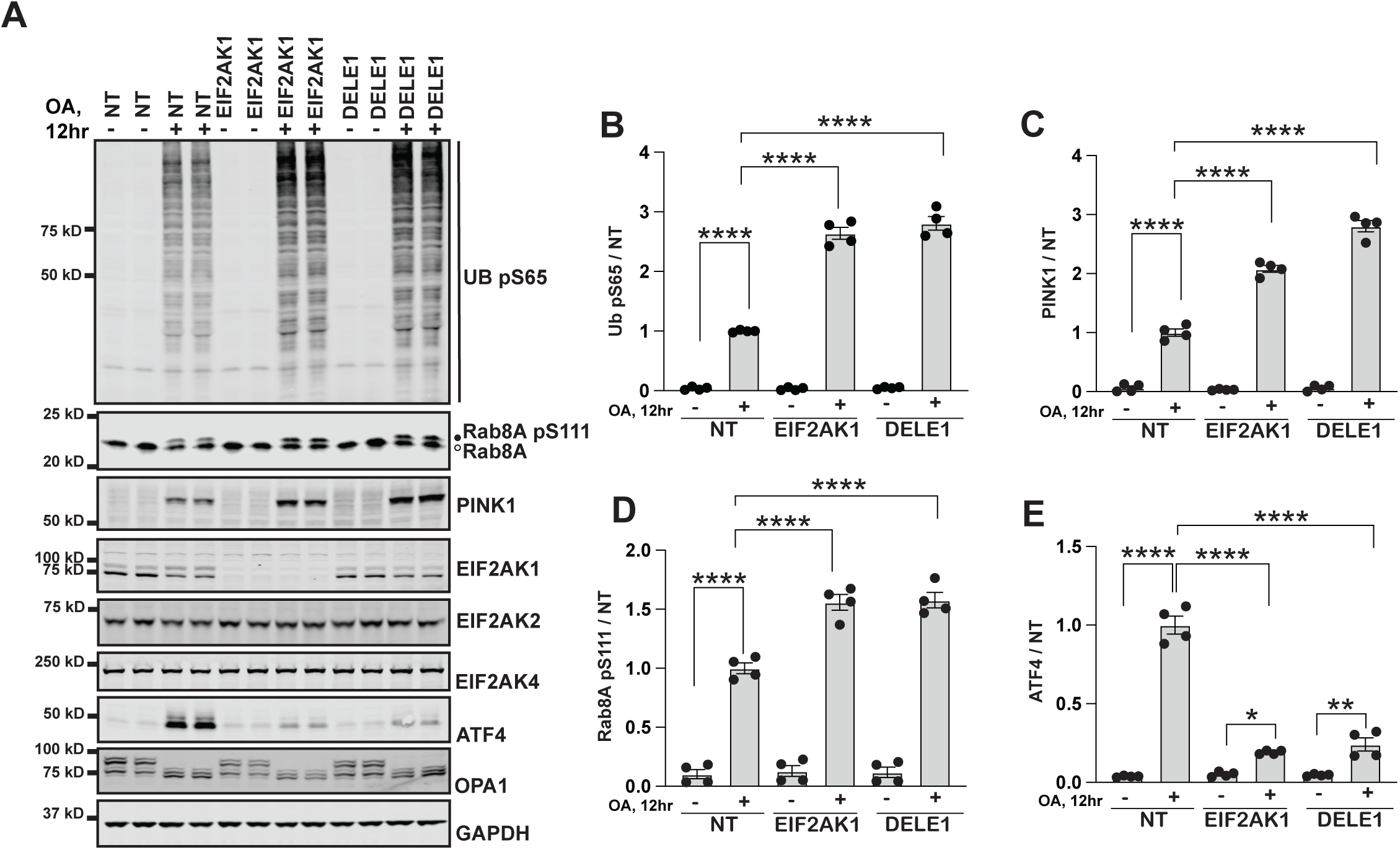
DELE1-EIF2AK1 signaling relay negatively regulates PINK1 stabilisation and activation. (**A**)HeLa cells either treated with non-targeting control siRNA pool (NT), siRNA pool targeting EIF2AK1 or, DELE1 for 72 hr were treated with OA in the last 12 hr. Immunoblot of Ub pS65, RAB8A, PINK1, EIF2AK1, EIF2AK2, EIF2AK4, ATF4, OPA1 and GAPDH. (**B**,**C**,**D** and **E**) Quantification for Ub pS65/GAPDH, PINK1/GAPDH, RAB8A pS111 and ATF4/GAPDH ratio for Fig. A, normalised to the ratio in NT (+OA) samples. Data information: (**B**,**C**,**D** and **E**) All data are mean ± SEM; Statistical significance is displayed as *P ≤ 0.05; **P ≤ 0.01; ***P ≤ 0.001; ****P ≤ 0.0001; ns, not significant. n = 4 technical replicates (2 biological replicates), one-way ANOVA, Tukey’s multiple comparisons test.

### Transcriptional and translational up-regulation of PINK1 in EIF2AK1 knockdown cells during mitochondrial damage-induced stress

Previous studies have shown that upon mitochondrial damage, PINK1 mRNA levels do not change and that PINK1 protein stabilisation is prevented by pre-treatment with the translation inhibitor, cycloheximide (CHX), suggesting a role for translation and new protein synthesis [31, 32]. Consistent with this we performed an RT-PCR timecourse experiment following OA treatment in HeLa cells and found that PINK1 mRNA levels do not significantly change upon mitochondrial damage (fig. S11A). This contrasted with a significant time-dependent increase in ATF4 mRNA levels following OA treatment (fig. S11B). We next measured PINK1 mRNA levels under conditions of siRNA-mediated knockdown of EIF2AK1. Using human TBP as an internal control, our RT-PCR results revealed no change in PINK1 mRNA levels in EIF2AK1 knockdown cells under basal conditions; however, we observed a significant upregulation of PINK1 mRNA in EIF2AK1 knockdown cells following OA-induced mitochondrial depolarisation (Fig. 4B). Consistent with this, co-treatment of cells with the transcriptional inhibitor, 5,6-dichlorobenzimidazole 1-β-D-ribofuranoside (DRB), completely prevented PINK1 mRNA up-regulation in OA-induced EIF2AK1 knockdown cells with only mild impact on basal PINK1 mRNA levels (Fig. 4B). The specificity of our RT-PCR assay was confirmed by complete loss of PINK1 mRNA in cells transfected with PINK1 siRNA (Fig. 4B). Further, under these RT-PCR assay conditions, the efficiency and specificity of EIF2AK1 knockdown was confirmed by prevention of OA-induced ATF4 mRNA up-regulation (fig. S11C), and complete abolishment of EIF2AK1 but not EIF2AK2 mRNA levels consistent with previous reports (fig. S11D, E). In parallel we assessed the impact of DRB co-treatment at the protein level in HeLa cells following EIF2AK1 knockdown and in agreement with the RT-PCR analysis, DRB prevented the OA-induced increase in PINK1 stabilisation and activation with only modest effects on PINK1 in non-targeting siRNA treated cells (Fig. 4D). We also observed similar results at the protein level using the transcriptional inhibitor, α-amanitin (fig. S11F). Furthermore, stabilization of PINK1 in EIF2AK1-silenced cells was inhibited by cycloheximide (CHX) treatment to a similar extent as non-targeting siRNA treated cells following OA treatment, confirming the role of protein translation (Fig. 4A, E and fig. S12 A, B). Overall these studies demonstrate that EIF2AK1 knockdown mediates transcriptional up-regulation and translation of PINK1 mRNA and in future work it will be interesting to uncover the transcriptional mechanism and how this mRNA undergoes translation on sites of mitochondrial damage.

**Fig. 4.**
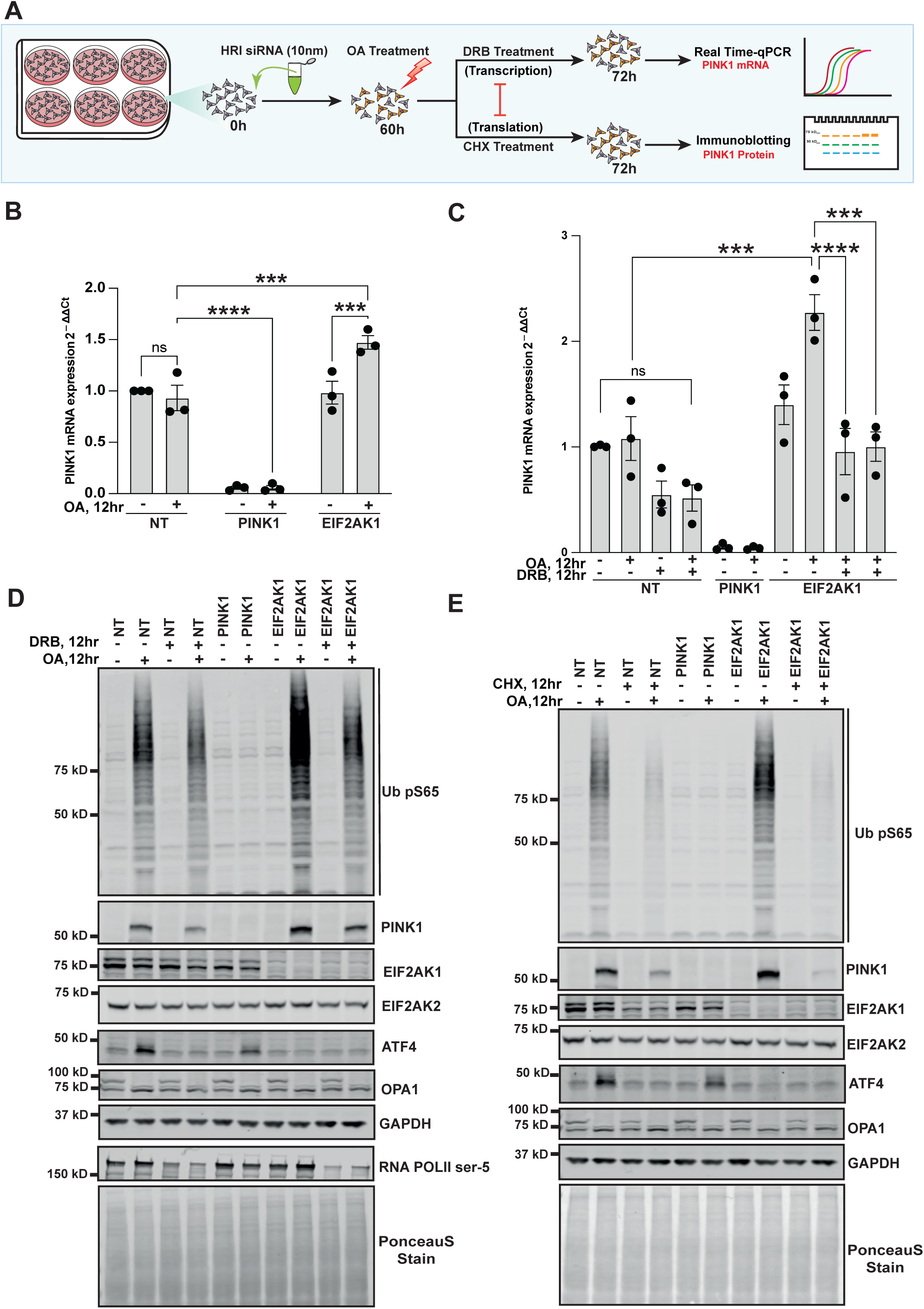
Transcriptional upregulation of PINK1 in EIF2AK1 knockdown cells following mitochondrial depolarisation. (**A**) Schematic of the siRNA workflow for studying transcriptional and translational regulation of PINK1. (**B**) HeLa cells either treated with non-targeting control siRNA pool (NT), siRNA pool targeting PINK1 or, EIF2AK1for 72 hr were treated with OA in the last 12 hr. Relative mRNA levels of PINK1 measured by RT-PCR in HeLa cells with TBP used as a reference gene. (**C**) as in B, Relative mRNA levels of PINK1 measured by RT-PCR in HeLa cells with and without co-treatment of transcription inhibitor DRB, using TBP as a reference gene. (**D**) Corresponding immunoblot analysis for remaing 2/3^rd^ of the samples used for RT PCR analysis in **C**. Immunoblot of Ub pS65, PINK1, EIF2AK1, EIF2AK2, ATF4, OPA1, GAPDH, RNA POLII phospho-S5 (DRB treatment control) and total protein as visualized by PonceauS staining. (**E**) HeLa cells treated with non-targeting control siRNA pool (NT), siRNA pool targeting PINK1 or, EIF2AK1for 72 hr were either treated with OA alone or, co-treated with translation inhibitor CHX, in the last 12 hr. Immunoblot of Ub pS65, PINK1, EIF2AK1, EIF2AK2, ATF4, OPA1, GAPDH and total protein as visualized by PonceauS staining. Data information: (**B** and **C**) All data are mean ± SEM; Statistical significance is displayed as *P ≤ 0.05; **P ≤ 0.01; ***P ≤ 0.001; ****P ≤ 0.0001; ns, not significant. n = 3 biological replicates, (**B**) two-way ANOVA, Uncorrected Fisher’s LSD multiple comparisons test and (**C**) one-way ANOVA, Tukey’s multiple comparisons test.

### Chemical inhibition of the ISR by ISRIB up-regulates PINK1 stabilization and activation in response to mitochondrial damage

The drug-like small molecule, Integrated Stress Response inhibitor (ISRIB), has been shown to antagonise eIF2α phosphorylation leading to inhibition of the ISR via promotion of translation and inhibition of ATF4 expression (Fig. 5A) [33, 34]. We therefore investigated the impact of ISRIB on mitochondrial damage-induced activation of PINK1 signaling. We initially undertook a timecourse of 300 nM ISRIB in HeLa cells treated with OA for 12 hr. This revealed that ISRIB significantly enhanced stabilisation and activation of PINK1 within 6 hr as measured by Ub pS65 and Rab8A pS111 (band-shift) and the effect of ISRIB was sustained up to 24 h (cells pre-treated with ISRIB for 12 h prior to addition of OA for 12 hr) (Fig. 5B–D). This was associated with ISRIB-induced inhibition of ATF4 following mitochondrial damage induced by OA (Fig. 5B, E). Furthermore, treatment with ISRIB alone did not affect PINK1 signaling in healthy cells, indicating that ISRIB’s effects are specific to cells undergoing mitochondrial stress and ISR pathway activation (fig. S13A). Consistent with these results in HeLa cells, we also treated ARPE-19 cells with ISRIB and observed an increase in total PINK1 levels and Ub pS65 following OA-induced mitochondrial damage compared to non-ISRIB treated cells (fig. S13B). Overall these chemical studies with ISRIB complement the above genetic findings, suggesting ISRIB-like small molecules as a potential therapeutic strategy to enhance PINK1-mitophagy.

**Fig. 5.**
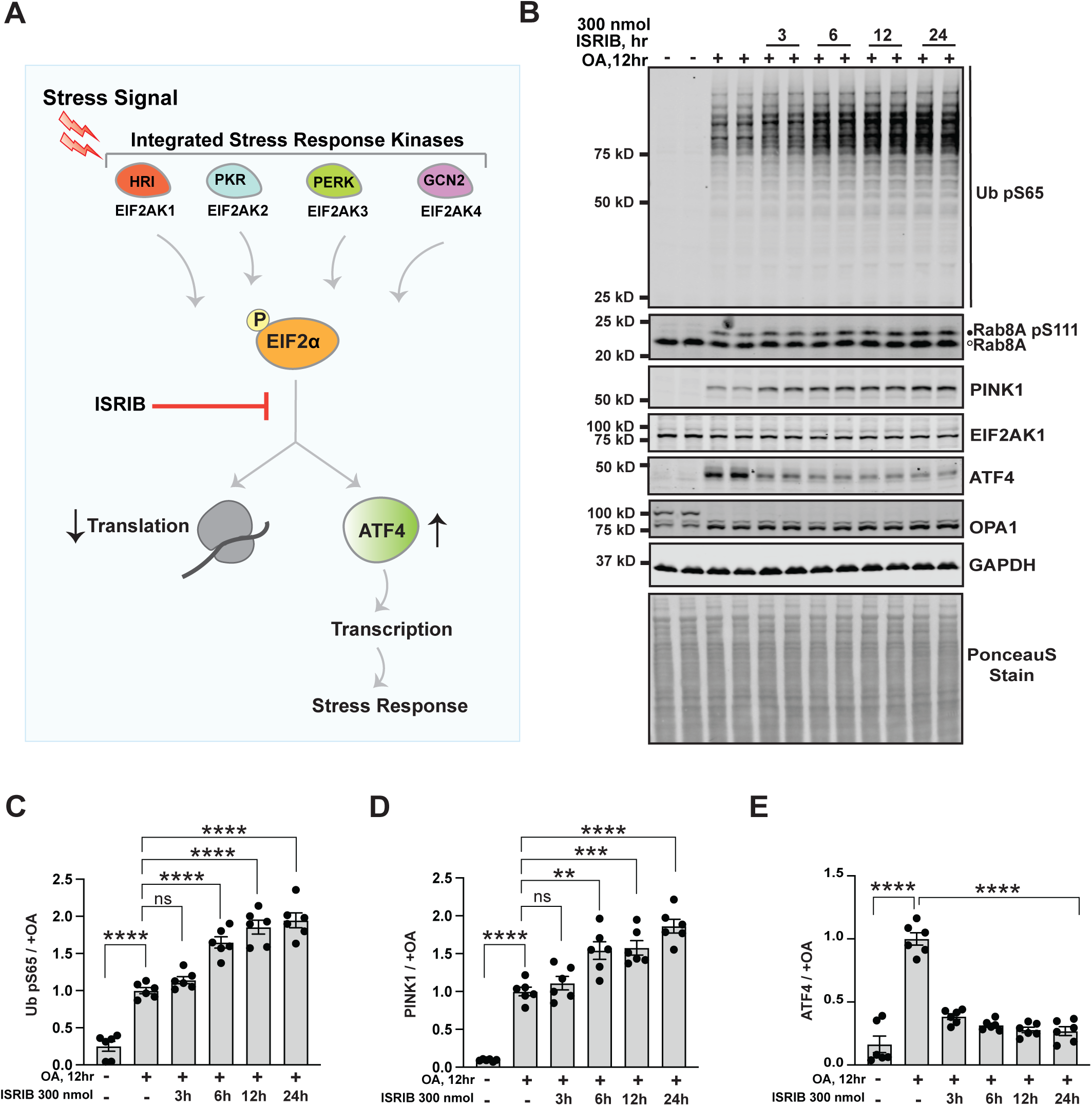
Chemical inhibition of the ISR by ISRIB enhances PINK1 stabilisation and activation. (**A**) Schematic depicting the two outputs of the ISR: reduction in bulk protein synthesis and translational induction of ATF4 and it’s target genes. ISRIB reverses these effects of the ISR by acting downstream of the ISR kinases phosphorylation of eIF2α. (**B**) HeLa cells seeded in a 10 cm dish, were co-treated with 300nmol of ISRIB for 3, 6, 12 and 24 hr and OA for 12 hr or, OA alone for 12 hr. Representative Immunoblot of Ub pS65, PINK1, Rab8A, EIF2AK1, OPA1, GAPDH, ATF4 and total protein as visualized by PonceauS staining.(**C**,**D** and **E**) Quantification for Ub pS65/GAPDH, PINK1/GAPDH and ATF4/GAPDH ratio for Fig. B, normalised to the ratio in OA alone samples using the Licor Image Studio software. Data information: (**C**,**D** and **E**) All data are mean ± SEM; Statistical significance is displayed as *P ≤ 0.05; **P ≤ 0.01; ***P ≤ 0.001; ****P ≤ 0.0001; ns, not significant. n = 6 technical replicates (3 biological replicates), one-way ANOVA, Tukey’s multiple comparisons test.

### Genetic or chemical inhibition of ISR enhances PINK1-dependent mitophagy

We next determined if the genetic knockdown of EIF2AK1 or chemical inhibition of ISR by ISRIB had an impact on PINK1-Parkin dependent mitophagy. To monitor mitophagy, we used the previously established and well-characterised *mito*-QC assay in Parkin overexpressing ARPE-19 cells [22, 23, 35]. This assay, utilising a stably expressed tandem mCherry-GFP tag attached to an OMM localisation peptide (derived from the protein FIS1), relies on a fluorescent colour change that occurs when mitochondria are delivered to lysosomes (referred to as mitolysosomes). Lysosomal acidity is sufficient to quench GFP, but not mCherry, and “red-only” mitolysosomes appear that can be easily quantified as a mitophagy readout. We observed that under basal conditions, siRNA knockdown of EIF2AK1 did not significantly increase mitophagy compared to NT-control ARPE-19 Parkin overexpressing cells (Fig. 6A-C). In contrast, loss of EIF2K1 resulted in a a significant increase in OA-induced mitolysosomes, indicative of increased mitophagy that is consistent with previous analyses of PINK1 activation (Fig. 6A-C). We next co-treated ARPE-19 Parkin overexpressing cells with or without OA and ISRIB. As with analysis of PINK1 signaling (fig. S13), we did not observe any significant effect of ISRIB on basal mitophagy (Fig. 6D-F). However, we observed a small but significant mitophagy increase in cells co-treated with OA and ISRIB compared to OA alone (Fig. 6D-F).

**Fig. 6.**
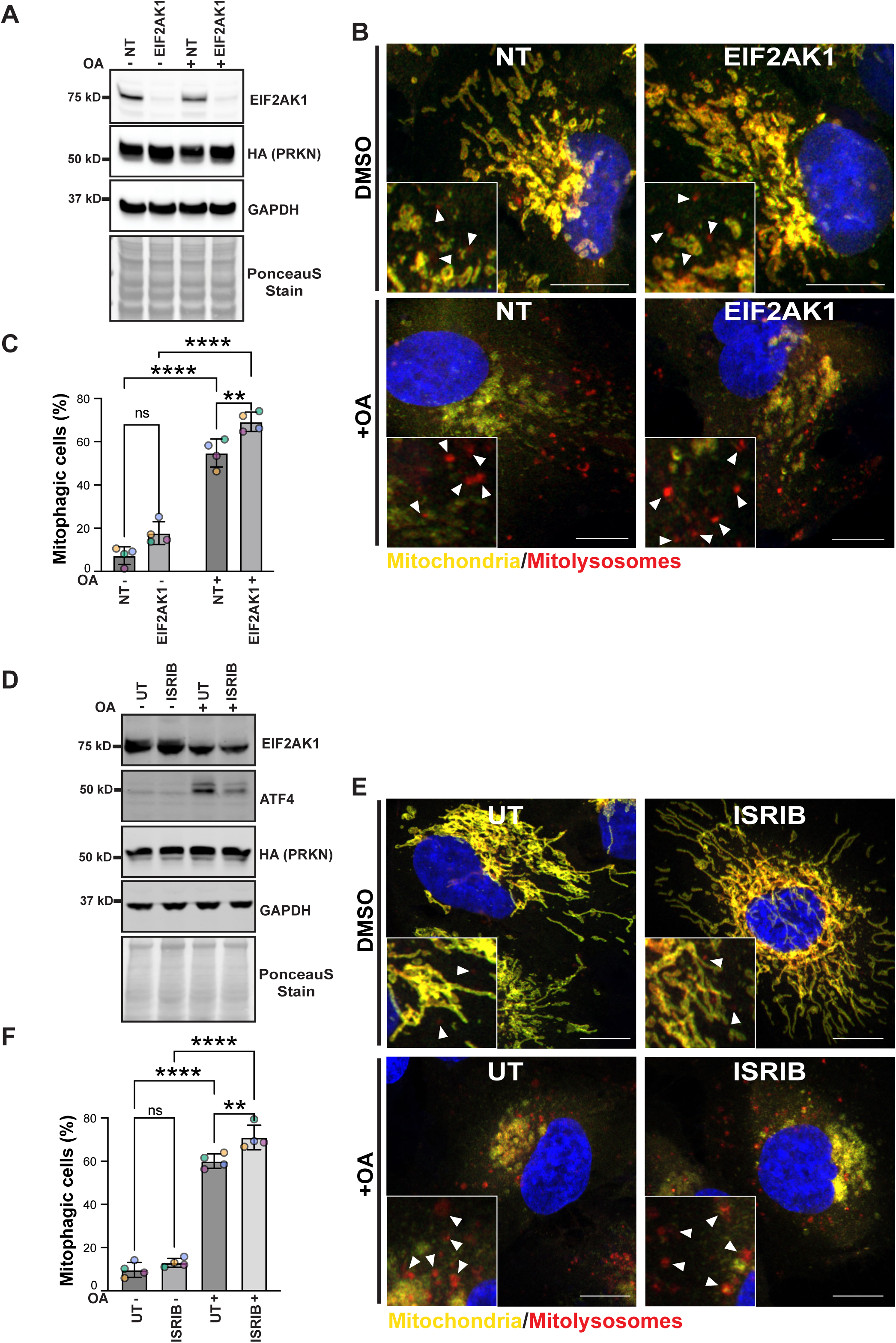
Genetic and chemical inhibition of of EIF2AK1-mediated ISR enhances PINK1-Parkin dependent mitophagy. (**A**,**B**) Representative immunoblots (**A**) or confocal images (**B**) of ARPE-19 cells stably expressing the mito-QC reporter and HA-Parkin, transfected with non-targeting siRNA (NT) or siRNA targeting EIF2AK1. 3 days post transfection, cells were treated with OA for 2 hr prior analysis. (**C**) Quantification of mitophagy shown in B from 4 independent experiments (n=4 biological replicates with >66 cells per replicate). (**D**,**E**) Representative immunoblots (**D**) or confocal images (**E**) of ARPE-19 cells stably expressing the mito-QC reporter and HA-Parkin, pre-treated 12 hr with 300 nM ISRIB and treated with OA for 2 hr prior analysis. F: Quantification of mitophagy shown in E from 4 independent experiments (n=4 biological replicates with >101 cells per replicate). (C) two-way ANOVA, Tukey’s multiple comparisons test and (F) one-way ANOVA, Tukey’s multiple comparisons test. Data information: Enlarged views are shown in the lower corners and arrowheads indicate examples of mitolysosomes. Nuclei were stained in blue (Hoechst). Scale bar: 10 µm. Overall data are mean ± s.d.; Statistical significance is displayed as *P ≤ 0.05; **P ≤ 0.01; ***P ≤ 0.001; ****P ≤ 0.0001; ns, not significant.

It was recently reported that EIF2AK1/HRI is a positive regulator of both PINK1-independent deferiprone (DFP)-mediated mitophagy and PINK1-Parkin-dependent OA-mediated mitophagy [36]. Deferiprone, also known as DFP, is an iron chelator that mimics hypoxic conditions through stabilisation of the oxygen-sensitive transcription factor HIF1α and has been described as the most potent mitophagy inducer in a PINK1-Parkin-independent manner [37]. We performed siRNA-mediated knockdown of EIF2AK1 in *mito*-QC expressing ARPE-19 cells treated with or without DFP in the presence or absence of BafilomycinA (BafA), a lysosomal inhibitor. Following DFP treatment, we observed significant reduction in the mitochondrial proteins HSP60, OMI, and COXIV in NT-targeted control cells that was prevented by co-treatment with BafA, indicating DFP-induced mitophagy (fig. S14 A-B). However, in EIF2AK1 depleted cells, mitochondrial proteins were similarly degraded upon DFP treatment and no significant difference was observed with the NT-targeted cells, demonstrating that mitophagy still occurs in EIF2AK1 knockdown cells (fig. S14 A-B). To validate these findings we measured mitophagy directly using Fluorescence Activated Cell Sorting (FACS)-based assay of the *mito*-QC reporter and did not detect any significant difference in mitophagy between EIF2AK1 knockdown and NT-targeted cells under our assay conditions (fig. S14 A-B). Overall our mitophagy analyses indicate that knockdown of EIF2AK1, or addition of ISRIB, enhances PINK1 mitophagy and that the ISR pathway acts as a specific negative regulator of OA-induced mitophagy but does not impact the DFP-induced mitophagy pathway.

## DISCUSSION

We have identified via an unbiased siRNA whole Ser/Thr kinome screen that the integrated stress response (ISR) kinase EIF2AK1/HRI acts as a negative regulator of PINK1-dependent signaling in cells. We show that under conditions of mitochondrial stress, genetic knockdown of EIF2AK1 or its stress-induced activator DELE1 inhibit the ISR leading to enhancement of PINK1 stabilisation and activation. Furthermore, we have discovered that the small molecule, ISRIB, a potent inhibitor of the ISR, (via antagonising the effects of eIF2α phosphorylation) enhances PINK1 activation and signaling. The physiological relevance of our findings is underscored by the demonstration that knockdown of EIF2AK1 or ISRIB stimulate PINK1-Parkin-dependent mitophagy induced by mitochondrial depolarisation. Overall our findings elaborate a model whereby the DELE1-EIF2AK1 relay pathway negatively interplays with the PINK1-Parkin mitophagy pathway (Fig. 7).

**Fig. 7.**
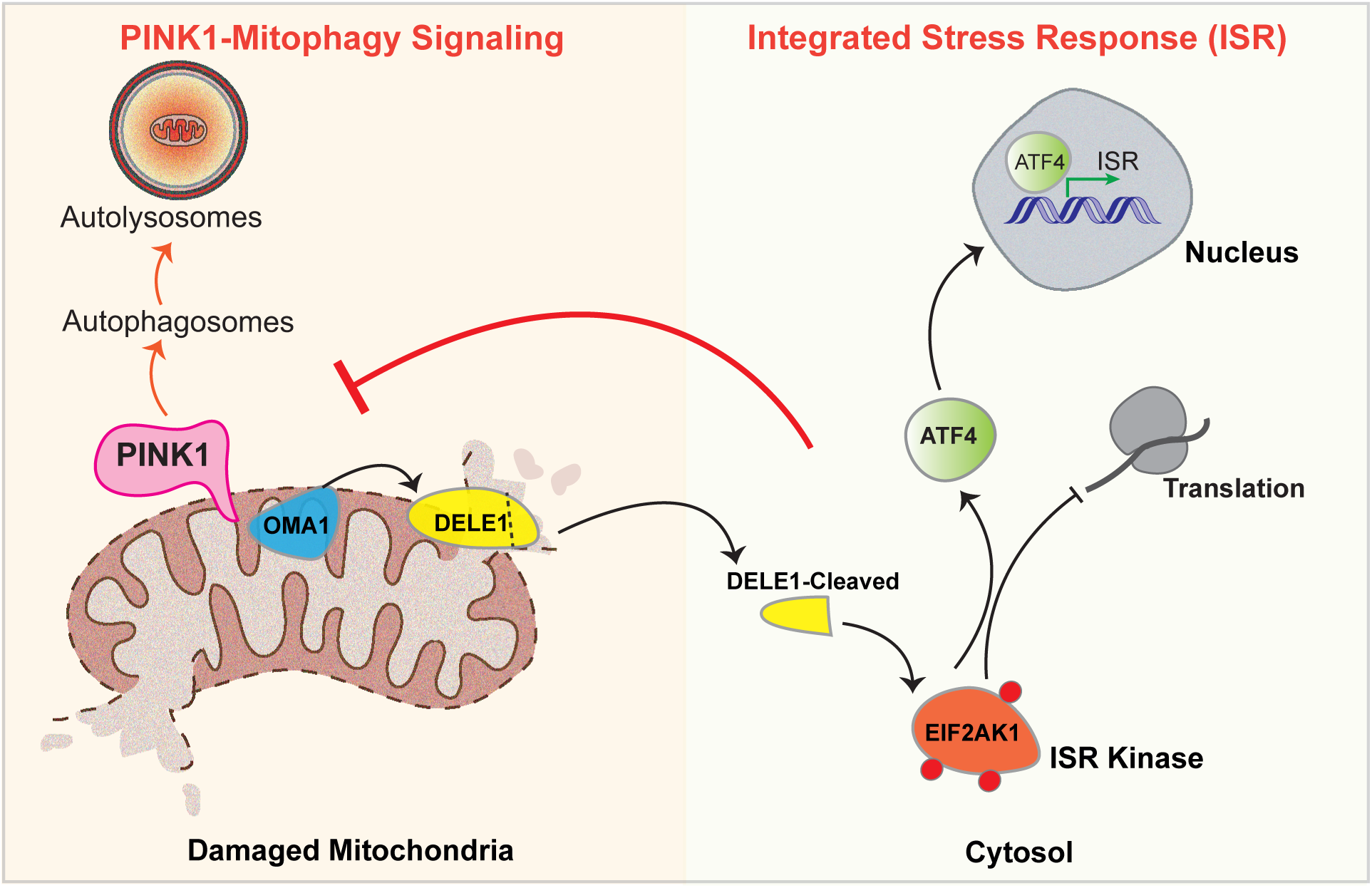
Model showing the negative interplay of DELE1-EIF2AK1 ISR pathway and PINK1 mitophagy signaling. Upon mitochondrial stress both PINK1 and OMA1 gets activated. OMA1 causes cleavage of DELE1 to a short form which accumulates in the cytosol, where it interacts with and activates the Integrated Stress Response (ISR) kinase EIF2AK1. EIF2AK1 activation leads to a reduction in global translation while allowing the translation of specific mRNAs including the transcription factor ATF4. Our findings suggest that silencing either DELE1 and EIF2AK1 branch of ISR or, chemical inhibition of ISR by ISRIB enhances the mitochondrial stress-induced activation and signaling of PINK1 and finally the autophagic removal of damaged mitochondria by Mitophagy.

Mitochondrial stress-induced activation of EIF2AK1 by DELE1 leads to phosphorylation of eukaryotic translation initiation factor 2α (eIF2α) resulting in the induction of the transcription factor ATF4 and global inhibition of mRNA translation and protein synthesis that mediate the ISR [18, 19]. In agreement with this, we observed time-dependent up-regulation of both ATF4 mRNA and protein levels following mitochondrial depolarisation and that was significantly reduced by either EIF2AK1 or DELE1 genetic knockdown. Initially identified as a transcriptional repressor of the cAMP element (CRE), ATF4 can act as both a transcriptional activator and inhibitor in response to various stressors and is a critical downstream effector modulating transcription of multiple genes including those regulating mitochondrial function and quality control [14–17, 27, 28]. The specific gene expression regulated by ATF4 has been attributed to combinations of several factors such as ATF4 heterodimerisation partners, post-translational and histone modifications surrounding target genes [14–17, 27, 28]. Previous studies indicated that PINK1 mRNA levels remain unchanged with mitochondrial stress, while the protein stabilizes rapidly unless pre-treated with the translation inhibitor CHX [31, 32]. Our results reveal that PINK1 mRNA transcription and translation is increased following mitochondrial depolarization under conditions of EIF2AK1 knockdown, and suggests that the ISR perhaps, via ATF4, may repress PINK1 transcription. Interestingly, the PINK1 gene promoter lacks typical TATA and CAAT boxes but, contains several putative and canonical regulatory sites for the transcription factors AP1F, NF-kB and CRE [38, 39]. However, how PINK1 is transcriptionally regulated by any stress is largely unexplored in the field and in future work it would be interesting to investigate the mechanism by which PINK1 mRNA is transcriptionally regulated by ATF4 under basal and mitochondrial stress conditions. Furthemore, mitochondrial proteins have been shown to be actively translated on the mitochondrial surface, and cryo-electron tomography-visualized ribosomes have been observed interacting with the TOM complex in both basal and mitochondrial depolarization states, facilitating co-translational import of select mitochondrial proteins although the effect of ISR stress in this context is not known [40, 41].

In a recent CRISPRi screen in human K562 cells, EIF2AK1/HRI was identified as a positive regulator of DFP-induced mitophagy [36]. The same group also reported that EIF2AK1 acted as a positive regulator for PINK1-Parkin mitophagy in HeLa cells that is contrary to our findings presented here [36]. At present we are unable to reconcile the differences between the studies although in our study we have been able to demonstrate that EIF2AK1 acts as a negative regulator of PINK1 mitophagic signaling in multiple human cell lines including HeLa, ARPE-19, SK-OV-3, and U2OS cells. Furthermore, under our assay conditions, the small molecule ISRIB also enhanced PINK1 stabilisation, activation and mitophagy in a similar way to what we observed with genetic knockdown of EIF2AK1, supporting a role for the ISR in negatively regulating PINK1 signaling following mitochondrial stress. The relatively smaller effect observed for EIF2AK1 knockdown or ISRIB on OA-induced mitophagy compared to OA-induced PINK1 activity signaling likely reflects the high level of mitophagy conferred by Parkin over-expression in this system via Parkin-mediated feed-forward amplification of downstream signaling. In future studies it would be interesting to determine whether EIF2AK1 regulates PINK1 in cells more relevant to PD including human iNeurons and primary mouse neurons that exhibit robust PINK1-Parkin dependent expression and signaling [42–44]. It has been reported in neurons that mRNA is co-transported with mitochondria along axons and locally translated to enable mitophagy in distal parts of the axon [45] and it would be interesting to investigate whether the levels of PINK1 mRNA in axons of neurons are regulated by ISR pathway activation induced by mitochondrial stress. It would also be important to evaluate whether EIF2AK1 and ISR regulate endogenous PINK1-Parkin dependent signaling and mitophagy *in vivo*. Using the *mito*-QC mouse reporter line, it has recently been reported that skeletal hindlimb muscle represents the tissue of choice for studying PINK1-dependent mitophagy *in vivo* with significant reduction of mitophagy in PINK1 knockout (KO) mice [46]. EIF2AK1 knockout mice have been reported to be viable and display mild alterations in erythrocytes but no gross morphological abnormalities in other tissues [26]. In future work, it would be interesting to cross EIF2AK1 knockout mice with the *mito*-QC mouse reporter line and determine whether EIF2AK1 loss leads to enhanced PINK1 mitophagy in skeletal muscle. It would also be interesting to treat wild-type and PINK1 KO mice with ISRIB or analogues to assess whether these can enhance PINK1-dependent mitophagy in skeletal muscle *in vivo*.

The original DELE1-HRI discovery studies demonstrated in HeLa cells that the mitochondrial protease OMA1 was critical for the cleavage of DELE1 with concomitant release of the C-terminal fragment of DELE1 into the cytosol [18, 19]. However, a recent study has reported that the mitochondrial protease HtrA2 can also mediate the cleavage of DELE1. In future work it will be interesting to determine the protease regulating DELE1 cleavage in cells that are physiologically relevant to PD including neurons [47]. Additionally, a recent study highlighted the critical role of the SIFI (Silencing Factor of the Integrated Stress Response) complex, an E3 ligase that degrades both the active form of EIF2AK1 and the cleaved DELE1 fragment along with accumulated unimported mitochondrial proteins during mitochondrial import stress [30]. This degradation process is vital for silencing the ISR and restoring the cellular equilibrium after stress. In our experiments, we also observed a reduction in endogenous EIF2AK1 levels following OA induced mitochondrial depolarisation, suggesting the involvement of the SIFI complex in this response and it would be interesting to investigate for potential crosstalk between the SIFI complex and PINK1-mediated mitophagic signaling in the future.

To date, the best-characterized negative regulator of PINK1/Parkin-induced mitophagy signaling is the deubiquitinase (DUB) USP30, discovered following an overexpression screen with a Flag-tagged human DUB cDNA library, with Phase I human trials of USP30 inhibitors underway in Parkinson’s patients (Mission Therapeutics) [48]. Our discovery that ISRIB enhances PINK1 stabilisation, activation, and mitophagy opens up novel therapeutic approaches to treat PD. ISRIB has been demonstrated to restore translational inhibition induced by ISR activation and to prevent gene expression induced by ATF4 by acting downstream of phosphorylated eIF2α [15, 33]. In mammalian cells exposed to ER stress inducers such as thapsigargin and tunicamycin, ISRIB effectively blocked the PERK-mediated induction of ATF4 [33]. Interestingly, ISRIB, was found to confer neuroprotection in select animal models *in vivo* [33, 34] and was recently reported to rescue neurodegeneration of Amyotrophic Lateral Sclerosis (ALS) gene VAPB iPSC-derived motor neurons [49]. Furthermore, two clinical drug analogues of ISRIB, ABBV-CLS-7262 (AbbVie/Calico) and DNL-343 (Denali Therapeutics) are currently undergoing evaluation in clinical trials of ALS [50]. Critically in our studies, ISRIB and EIF2AK1 knockdown did not impact PINK1 signaling in healthy basal cells and was specific to cells undergoing mitochondrial stress suggesting that therapeutic approaches to target the ISR such as ISRIB or inhibition of EIF2AK1 activation by DELE1 may be associated with fewer side-effects.

Overall, our current analysis strongly indicates that mitochondrial stress-induced activation of the ISR, mediated by the EIF2AK1 / HRI kinase, is a negative regulator of PINK-dependent mitophagic signaling. These insights offer new potential therapeutic strategies against PD.

## MATERIALS AND METHODS

### Materials

cDNA constructs for mammalian tissue culture were amplified in *Escherichia coli* (*E. coli*) DH5*α* cells and purified using a NucleoBond Xtra Midi kit (no. 740410.50; Macherey-Nagel). All DNA constructs were verified by DNA sequencing, which was performed by the Sequencing Service, School of Life Sciences, University of Dundee, using DYEnamic ET terminator chemistry (Amersham Biosciences) on Applied Biosystems automated DNA sequencers. DNA for bacterial protein expression was transformed into *E. coli* BL21 DE3 RIL (codon plus) cells (Stratagene). All cDNA plasmids, CRISPR gRNAs, antibodies and recombinant proteins generated for this study are available via our reagents website https://mrcppureagents.dundee.ac.uk/.

### Reagents

Antimycin A (A8674), Oligomycin (75351), Cycloheximide (CHX) (C7698), 5,6-Dichlorobenzimidazole 1-β-D-ribofuranoside (DRB) (D1916), ISRIB (SML0843) and Deferiprone (DFP) (379409), were purchased from Sigma-Aldrich. alpha-Amanitin, (4025) was purchased from Tocris/BioTechne. Bafilomycin A1 ATPase inhibitor (BafA) (BML-CM110) was purchased from Enzo Life Sciences.

### Antibodies for biochemical studies

The following primary antibodies were used: Ubiquitin phospho-Ser65 (Cell Signaling Technology (CST Cat# 62802)), OPA1 (BD BIOSCIENCES Cat# 612607), Rab8A (Abcam Cat# Ab241061), Rab8A phospho-Ser111 (Abcam Cat# Ab267493), PINK1 (Novus Cat# BC100-494, CST Cat# 6946T), PINK1 (in-house generated by Dundee Cell Products (DCP)), GAPDH (Santa Cruz Cat# sc-32233), EIF2AK2 (Abcam Cat# Ab184257), EIF2AK3(Abcam Cat# Ab229912), EIF2AK4(Abcam Cat# Ab134053), ATF4 (CST Cat# 11815S), OMA1 (Santa Cruz Cat# sc-515788), TRIM28 (Anti-KAP1) (Abcam Cat# Ab109287), RPS6KB2 (Abcam Cat# Ab184551), BCR (Abcam Cat# Ab233709), BRD2 (Abcam Cat# Ab139690), MAP3K7 (Abcam Cat# Ab109526), CDC7 (Abcam Cat# Ab229187), ROCK1 (Abcam Cat# Ab134181), EEF2K (Abcam Cat# Ab45168), CAMK1D (Abcam Cat# Ab172618), GRK2 (Abcam Cat# Ab227825), CNSK2A1 (Abcam Cat# Ab70774), CSNK1A1 (Abcam Cat# Ab206652), GSK3B (Abcam Cat# Ab32391), MAPK3 (Abcam Cat# Ab32537), PBK/SPK (Abcam Cat# Ab236872), MAP3K5 (Abcam Cat# Ab45178), MAP2K2 (Abcam Cat# Ab32517), MAP2K6 (Abcam Cat# Ab33866), MAP2K7 (Abcam Cat# Ab52618), HA (Invitrogen Cat# 26183), Anti-RNA polymerase II CTD repeat YSPTSPS(phosphor S5) (Abcam Cat# Ab817), HIF1α (R&D system Cat# MAB1536), HSP60 (CST Cat# 4870S), COXIV (CST Cat# 4850S), anti-LC3 A/B (CST Cat# 4108S), VINCULIN (Abcam Cat# Ab129002). The following polyclonal antibodies were produced by the MRCPPU Reagents and Services at the University of Dundee in sheep: EIF2AK1 (S085D), OMI (S802C).

### siRNA screens and follow-up experiments

The siRNA screens were performed using a human siRNA library (Horizon Dharmacon) designed to target 428 ser/Thr kinases. The list of target siRNA pools and their oligonucleotide sequences are listed in Table S2. 1.5 ml of Hela cells were seeded in 6-well plates at 37,000 cells/ml and transfected using 10nm of siRNA and 1.5 μl of the Lipofectamine™ RNAiMAX transfection Reagent (Thermo Fisher Scientific) per well. Cells were cultured for 52 hr after which were stimulated with Oligomycin (1 μM) and Antimycin (10 μM) (OA). After further 20 hr cells were lysed in 50 μl of Lysis buffer, centrifuged at 17,000 g for 15 min at 4°C, quantified by Bradford Assay (Thermo Scientific) and subjected to immunoblot analysis. Each siRNA screening experiment also included untreated cells, positive control PINK1 targeting siRNA, and a negative control siRNA (non-targeting siRNA, NT). The siRNA studies for further validation of EIF2AK1 with independent pooled siRNA for EIF2AK1, EIF2AK2, EIF2AK3 and EIF2AK4 as well as single siRNAs for EIF2AK1 (Dharmacon, Table S2) were also performed in four different cell lines (HeLa, SKOV3, U2OS, ARPE-19) as above. However, in the further validation studies the Cells were cultured for 60 hr and then stimulated with Oligomycin (1 μM) and Antimycin (10 μM) (OA). After further 12 hr cells were lysed in 50 μl of Lysis buffer, centrifuged at 17,000 g for 15 min at 4°C, quantified by Coomassie (Bradford) Protein Assay Kit (WTS) and subjected to immunoblot analysis. The siRNA studies for targeting DELE1 (Horizon Dharmacon, Table S2) alongside EIF2AK1 were also performed in a similar manner.

### Generation of EIF2AK1 CRISPR/Cas9 knockout cells

A full transcript map of the EIF2AK1 locus was constructed by combining data from both NCBI (NC_000007.14 (6022247..6059175, complement) and Ensembl (ENSG00000086232). Knockout (KO) guides were chosen to target as far upstream as possible within an exon common to all published and predicted variants to ensure complete disruption of all possible transcripts following CRISPR targeting. Three pairs of CRISPR/Cas9 guides were designed: A pair targeting exon 2; G1 single guide RNA targeting exon 2; and G2 single guide RNA targeting exon 2 of the EIF2AK1 gene and this would be predicted to abolish expression of full-length EIF2AK1 protein. (fig. S9A-D). Complementary oligonucleotides were designed and annealed to generate dsDNA inserts with overhangs compatible to BbsI digested destination vectors. The sense guide insert was subsequently cloned into BbsI digested pBABED P U6 (DU48788, University of Dundee) and the antisense cloned into BbsI digested pX335 (Addgene #42335), yielding clones DU69746 and DU69747 respectively.

HeLa cells were co-transfected with 1μg of each plasmid using PEI in a 10cm dish. Following 24 hr of recovery and a further 48 hr of puromycin selection (1μg/ml). The cell pool was subsequently analysed for EIF2AK1 depletion by immunoblotting then single cell sorted via FACS. Following recovery, individual clones were analysed for EIF2AK1 depletion by immunoblotting (fig. S9E) and the promising clones A2 and A3 further verified by PCR, shotgun cloning and sequencing. Briefly, genomic DNA was isolated and a 1.8 Kb region containing exon 2 was amplified by PCR (forward primer: CACGGCATCTTTCTGCTGATCC; reverse primer: TCCAATTTTTGTATACCAGACGCTTTCC) (fig. S9B,C). The resulting PCR products were subcloned into the holding vector pSC-B (StrataClone Blunt PCR Cloning Kit, Agilent Technologies) and 16 colonies (white) picked for each of the clonal lines. Plasmid DNAs were isolated and cut with EcoRI to verify insert size and 14 positive samples for each line sent for sequencing with primers M13F and M13R. Indel formation often leads to a wide range of variations between targeted alleles leading to a variety of variations between targeted alleles thus direct sequencing of amplified amplicon pools yields poor-quality data around the CRISPR target site(s). We have found in practice that analysis of >8-10 shotgun clones from a given clonal line is sufficient to verify the allelic population with precision, for hyper and hypotriploid lines more sequencing reads may be required to be confident that all alleles are accounted for. Sequencing of the A2 and A3 clones showed 3 copies of the chromosome in each case and each indel was confirmed to lead to frameshift and premature termination of EIF2AK1 further corroborating the western data and confirming successful KO.

### Cell culture and transfection

HeLa, U2OS and SK-OV-3 cells were routinely cultured in standard DMEM (Dulbecco’s modified Eagle’s medium) supplemented with 10% (v/v) FBS, 2 mM L-glutamine, 100 U/mL penicillin and 0.1 mg/mL streptomycin. ARPE-19 cells (ATCC, CRL-2302) were routinely cultured in 1:1 DMEM:F-12 media supplemented with 10% (v/v) FBS, 2 mM L-glutamine, 100 U/mL penicillin and 0.1 mg/mL streptomycin. The cells were passaged by washing the cells (80-90% confluency) with PBS followed by incubation with Trypsin/EDTA. To inactivate Trypsin/EDTA (1:1) pre-warmed cell culture medium was added after 5 min and cells centrifugated at 1200 rpm for 3 min. The cell number in the suspension was counted by an automated cell counter and seeded into new culture dishes at required densities. All cells were cultured at 37°C, 5% CO_2_ in a humidified incubator and routinely tested for mycoplasma. Where indicated, cells were treated with Oligomycin (1 μM) /Antimycin A (10 μM) (OA) for mitochondrial depolarisation and 300 nM ISRIB as specified for chemical inhibition of the ISR. For iron chelation DFP was used at 1 mM concentration and inhibition of lysosomal degradation was achieved by Bafilomycin A (BafA) treatment at 50 nM. Inhibition of transcription was achieved by treatment with 5,6-dichlorobenzimidazole (DRB) at 50 μM, alpha-Amanitin at 5ug/ml and translation by treatment with cycloheximide (CHX) at 1μg/ml for 12h.

### Whole-cell lysate preparation

Cells were lysed in an ice-cold lysis buffer containing 50 mM Tris-HCl, pH 7.45, 150mM NaCl, 1% (by vol) Triton X-100, 5mM MgCl_2_, 1 mM sodium orthovanadate, 50 mM NaF, 10 mM 2-glycerophosphate, 5 mM sodium pyrophosphate, 0.5 ug/ml Microcystin LR (Enzo) and complete EDTA-free protease inhibitor cocktail (Roche) with freshly added 1x phosphatase inhibitor cocktail (Sigma-Aldrich) and 2ul/ml Benzonase (SIGMA). Lysates were clarified by centrifugation at 17, 000g at 4°C for 15 min, and supernatants were quantified by Bradford Assay (Thermo Scientific) and Bicinchoninic Acid Assay (Thermo Scientific).

### Immunoblotting

Samples were subjected to SDS-PAGE (Tris-glycine 12% gels, Bis-Tris 4-12% gels; Novex) and transferred onto nitrocellulose membranes. Membranes were blocked for 1h at room temperature with 5 % non-fat milk or 5% BSA in Tris buffered saline (TBST; 50 mM Tris-HCl and 150 mM NaCl, pH 7.5) containing 0.1 % Tween-20 and incubated at 4°C overnight with the indicated antibodies, diluted in 5% bovine serum albumin (BSA) or, 5 % non-fat milk. Highly cross-absorbed H+L secondary antibodies (Life Technologies) conjugated to (IRDye® 800CW or IRDye® 680RD Infrared Dyes) were used at 1:10000 in TBST for 1h and the membrane was washed once with TBS then imaged using the OdysseyClx Western Blot imaging system.

### Protein purification from *E. coli*

Full-length wild-type EIF2AK1 was expressed in *E. coli* as N-terminal GST fusion protein and purified as described previously for GST proteins [51]. Briefly, BL21 Codon+ transformed cells were grown at 37 °C to an OD_600_ of 0.3, then shifted to 16 °C and induced with 250 μM IPTG (isopropyl β-D-thiogalactoside) at OD_600_ of 0.5. Cells were induced with 250 μM IPTG at OD 0.6 and were further grown at 16 °C for 16 h. Cells were pelleted at 4000 r.p.m., and then lysed by sonication in lysis buffer containing 50 mM Tris-HCl (pH 7.5), 150 mM NaCl, 0.1 mM EGTA, 0.1 % (v/v) 2-mercaptoethanol, 270 mM sucrose. Lysates were clarified by centrifugation at 30 000 x g for 30 min at 4 °C followed by incubation with 1 ml of GSH Agarose resin for 1.5 h at 4 °C. The resin was washed thoroughly in wash buffer containing 50 mM Tris-HCl (pH 7.5), 200 mM NaCl, 0.5 mM TCEP, and 10% glycerol, and the protein was eluted by incubation with wash buffer containing 10 mM glutathione for 1 hr at 5° to 7°C. The eluted supernatant was dialyzed against wash buffer at 5° to 7°C overnight and concentrated, and the final sample was flash-frozen.

### Quantitative RT-PCR

HeLa cells seeded in 10 cm dishes were washed and collected in sterile PBS. Samples were divided such that 2/3^rd^ cells were used for immunoblot analysis as described above and the remaining 1/3^rd^ were snap frozen for total RNA isolation. RNA was extracted for each sample using the PureLinkTM RNA Mini Kit (Invitrogen by Thermo Fisher Scientific, #12183025). cDNA synthesis was achieved using the PrimeScriptRT reagent Kit with gDNA Eraser (Takara, #RR047A) with 1 μg of RNA as template and following manufacturer’s instructions. The obtained cDNA was used as a template for qPCR with TB Green® Premix Ex TaqTM II (Tli TNaseJ Plus) (Takara, #RR820L) and the following primers (Sigma):

PINK1: 5′-AGACGCTTGCAGGGCTTTC-3′ (F) & 5′-GGCAATGTAGGCATGGTGG-3′ (R);

β-actin:5′-AGAAGGATTCCTATGTGGGCG-3′(F) & 5′-CATGTCGTCCCAGTTGGTGAC-3′ (R);

TBP:5’-TGTATCCACAGTGAATCTTGGTTG-3’ (F), & 5’-GGTTCGTGGCTCTCTTATCCTC-3’ (R);

ATF4:5′-CCAACAACAGCAAGGAGGAT-3′(F) & 5′-GGGGCAAAGAGATCACAAGT-3′ (R);

HRI: 5’-ACCCCGAATATGACGAATCTGA-3’ (F) & 5’-CAAGTGCTCCAGCAAAGAAAC-3’ (R);

PKR: 5’-GCCGCTAAACTTGCATATCTTCA-3’(F) & 5’-TCACACGTAGTAGCAAAAGAACC-3’ (R). Real-time quantitative PCR (qPCR) was performed in triplicates on thermocycler CFX Opus 384 (Bio-Rad Laboratories) operated by CFX Maestro software (Bio-Rad) using cycling protocol of 30 sec at 95°C followed by 40 cycles of 5 sec at 95°C and 60 sec at 60°C, and followed by melting curve from 65°C to 95°C. The fold change in expression was calculated using the 2(-Delta Delta C(T)) method [52]. Data were analysed in Microsoft Excel and GraphPad Prism version 10.0.3. using one-way ANOVA (Multiple comparisons).

### Mitophagy assay

Cells stably expressing mito-QC mitophagy reporter system (mCherry-GFP-FIS1^101–152^) and HA-Parkin, were seeded onto sterile glass coverslips in 24-well dishes. After treatment, coverslips were washed twice with PBS, fixed with 3.7% (w/v) formaldehyde, 200 mM HEPES pH 7.0 for 10 min and washed twice with PBS. After nuclei counterstaining with 1 μg/ml Hoechst-33258 dye, slides were washed and mounted in ProLong Gold (Invitrogen). For quantification, images were taken with Nikon Eclipse Ti widefield microscope (Plan Apo Lambda 60× Oil Ph3 DM). All the images were processed with FIJI v1.52n software (ImageJ, NIH). Quantification of mitophagy (red-only dots) was performed from four independent experiments counting over 66 cells per condition. Images were processed with the mito-QC Counter plugin as previously described [22]. Note, we observed some variation between cells and replicats. To keep quantitation consistent, we set a threshold above which the cells were considered as mitophagic based on the untreated WT condition. As quantitation was automated, initial blinding to sample ID was not performed.

### Flow cytometry analysis

ARPE-19 cells were seeded in a 6 cm dish. After treatment, cells were washed with PBS, trypsinized for 5 min and centrifuged 3 min at 1,200 rpm. The pellet of cells was resuspended in 250 μl of PBS and 2 ml of 3.7% (w/v) formaldehyde, 200 mM HEPES pH 7.0 were added. After 30 min at RT, 3 ml of PBS was added before centrifugation 5 min at 1,200 rpm. Finally, the pellet of cells was resuspended in 1% FCS in PBS and directly analysed by flow cytometry. For each independent experiment, at least 5 × 10^4^ cells were acquired on LSRFortessa cell analyser. Based on FCS and SSC profiles, living cells were gated. As negative control, cells expressing any mitophagy reporter can be used. To quantify the percentage of cells underdoing mitophagy, the ratio GFP/mCherry was analysed. The gate used for the nontreated condition or control cells was applied to all the other conditions. The value used for this was based on quantitation of microscopy data from mito-QC cells that showed around 7% (mito-QC) of cells had red-only puncta above the value of the mean.

### Statistical analyses

All statistical analyses were performed in GraphPad Prism version 10.0.3. Representative results of at least two or three independent experiments (biological replicates) are shown in all panels as specified. For immunoblot quantifications, level of each protein were normalised to GAPDH or, VINCULIN and expressed as fold change. Statistical significance was determined either by ordinary one-way ANOVA or by ordinary two-way ANOVA with the appropriate multiple correction test. P-values are indicated as *P < 0.05, **P < 0.01, ***P < 0.001 and ****P < 0.0001. ns: P > 0.05. The details of each statistical test, n numbers and graph used are specified in the relative figure legends.

### HEK-293 transfection

HEK293 cells were transiently transfected with 5μg of each of C-terminal HA-tagged EIF2AK1 wild-type (WT) or Kinase-inactive (KI) or wild-type (WT) EIF2AK4 plasmid for 24 hr using PEI in a 10cm dish. 24 hr post-transfection, cells were lysed and immunoblotted with EIF2AK1 and HA antibodies (fig. S5 B, C) and membranes were analysed using the OdysseyClx Western Blot imaging system.

## ACKNOWLEDGEMENTS

We thank Brett Benedetti and Shalini Padmanabhan at the Michael J Fox Foundation for helpful discussions throughout the project. We thank Anne Bertolotti at the MRC-LMB for valuable discussions on the data. We thank Richard Youle for providing the S-HeLa PINK1 knockout cells. We are grateful to the sequencing service (School of Life Sciences, University of Dundee); James Hastie for expression and generation of EIF2AK1 recombinant protein (MRC PPU); the MRC PPU tissue culture team (co-ordinated by Edwin Allen) and MRC PPU Reagents and Services antibody teams (co-ordinated by James Hastie). This work was supported by the Michael J. Fox Foundation (M.M.K.M.), a Wellcome Trust Senior Research Fellowship in Clinical Science (210753/Z/18/Z to M.M.K.M.); EMBO YIP Award (M.M.K.M.) and the Medical Research Council (MC UU 00018/8 to A.R.; MC_UU_00018/2 to I.G.G). A.K. was funded as part of the University of Dundee School of Life Sciences Summer School Program.

