## Supplementary File for "Kinome screening identifies integrated stress response kinase EIF2AK1 / HRI as a negative regulator of PINK1 mitophagy signaling"

#### SUPPLEMENTARY FIGURE LEGENDS

**Fig. S1. Immunoblot analysis of siRNA Ser/Thr kinome screen for regulators of PINK1-mediated Rab8A Ser111 phosphorylation.**
(**A-G**) HeLa cells transfected with the indicated siRNA pools of 108 set of Ser/Thr Kinases (Dharmacon) for 72 hr and treated with OA in the last 20 hr. Cells were lysed and immunoblotted for the indicated antibodies: Rab8A pS111, Rab8A, PINK1, GAPDH, OPA1, BCR, TRIM28, RPS6KB2, BRD2. All the immunoblots were developed using the LI-COR Odyssey CLx Western Blot imaging system.

**Fig. S2. Immunoblot analysis of siRNA Ser/Thr kinome screen for regulators of PINK1-mediated Rab8A Ser111 phosphorylation.**
(**A-G**) HeLa cells transfected with the indicated siRNA pools of 123 set of Ser/Thr Kinases (Dharmacon) for 72 hr and treated with OA in the last 20 hr. Cells were lysed and immunoblotted for the indicated antibodies: Rab8A pS111, Rab8A, PINK1, GAPDH, OPA1, MAP3K7, CDC7, ROCK1, EEF2K, CAMK1D, GRK2. All the immunoblots were developed using the LI-COR Odyssey CLx Western Blot imaging system.

**Fig. S3. Immunoblot analysis of siRNA Ser/Thr kinome screen for regulators of PINK1-mediated Rab8A Ser111 phosphorylation.**
(**A-G**) HeLa cells transfected with the indicated siRNA pools of 126 set of Ser/Thr Kinases (Dharmacon) for 72 hr and treated with OA in the last 20 hr. Cells were lysed and immunoblotted for the indicated antibodies: Rab8A pS111, Rab8A, PINK1, GAPDH, OPA1, CSNK1A1, CSNK2A1, MAPK3, EIF2AK4, PBK/SPK. All the immunoblots were developed using the LI-COR Odyssey CLx Western Blot imaging system.

**Fig. S4. Immunoblot analysis of siRNA Ser/Thr kinome screen for regulators of PINK1-mediated Rab8A Ser111 phosphorylation.**
(**A-G**) HeLa cells transfected with the indicated siRNA pools of 71 set of Ser/Thr Kinases (Dharmacon) for 72 hr and treated with OA in the last 20 hr. Cells were lysed and immunoblotted for the indicated antibodies: Rab8A pS111, Rab8A, PINK1, GAPDH, OPA1, MAP3K5, MAP2K2, MAP2K6, MAP2K7. All the immunoblots were developed using the LI-COR Odyssey CLx Western Blot imaging system.

**Fig. S5. Expression of recombinant full-length GST-EIF2AK1 protein for generation and validation of polyclonal anti-EIF2AK1 antibody.**
(**A**) SDS-PAGE profile of N-terminal GST-tagged EIF2AK1 protein expressed and purified from *E. coli*. (**B**) HEK-293 cells transfected with C-terminal HA-tagged EIF2AK1 wild-type (WT) or Kinase-inactive (KI) and wild-type (WT) EIF2AK4 for 24 hr. Cells were lysed and immunoblotted using total human EIF2AK1 polyclonal antibody generated against full-length human N-terminal GST-tagged EIF2AK1. (**C**) As in B, Immunoblotting was done using HA antibody for detection of levels of EIF2AK1 and EIF2AK4.

**Fig. S6. Validation of specificity of siRNA-mediated knockdown of EIF2AK1.**
(**A**) HeLa cells were transfected with independent pooled siRNA (Sigma) as well as single siRNAs 1, 2, 3 and 4 (Dharmacon) for EIF2AK1 for 72 hr and treated with OA in the last 12 hr. Cells were lysed and immunoblotted for Ub pS65, PINK1, Rab8A, EIF2AK1, OPA1, GAPDH and immunoblots were developed using the LI-COR Odyssey CLx Western Blot imaging system.

**Fig. S7. Analysis of EIF2AK1 knockdown on PINK1 stabilisation and activation across a panel of human cell lines.**
(**A**) SK-OV-3 (**B**) ARPE-19 and (**C**) U2OS cells, either untreated (UN) or, treated with non-targeting control siRNA pool (NT), siRNA pool targeting PINK1, EIF2AK1, EIFF2AK2, EIF2AK3, EIF2AK4, for 72 hr were treated with OA in the last 12 hr. Cells were lysed and immunoblotted for Ub pS65, RAB8A, PINK1, EIF2AK1, EIFF2AK2, EIF2AK3, EIF2AK4, OPA1, GAPDH and ATF4 and the immunoblots were developed using the LI-COR Odyssey CLx Western Blot imaging system.

**Fig. S8. Quantitation of PINK1 stabilisation and activation in EIF2AK1 knockout CRISPR pool lysates.** (**A**, **B** and **C**) Quantification for Ub pS65/GAPDH, PINK1/GAPDH and Rab8A pS111/total Rab8A ratio normalised to the ratio in WT+OA samples for Fig. 2F using the Licor Image Studio software.
Data information: (**A**, **B** and **C**) All data are mean ± SEM; Statistical significance is displayed as $^{*}P \leq 0.05$; $^{**}P \leq 0.01$; $^{***}P \leq 0.001$; $^{****}P \leq 0.0001$; ns, not significant. n = 3 technical replicates (2 biological replicates), one-way ANOVA, Tukey's multiple comparisons test.

**Fig. S9. Generation, characterization and validation of PINK1 stabilisation and activation in two independent EIF2AK1 knockout clones.**
(**A**) The sequence of EIF2AK1 in wild-type (WT) HeLa cells at exon 2 is shown and the CRISPR A pair guides used to generate EIF2AK1 Knockout (KO) is highlighted as black arrows. (**B**) The flanking primer position used to confirm successful KO generation is highlighted (**C**) PCR analysis of the wild-type, EIF2AK1 knockout clone A2 and A3 using exon2 flanked forward and reverse primer. PCR products were further cloned and sequenced to confirm homozygous knockout alleles. (**D**) Reference exon 2 translated DNA sequence of wild-type (WT) and sequencing confirming alterations in the HeLa EIF2AK1 KO clone A2 and A3 cell lines has been highlighted. These nucleotide deletion and insertion changes in exon 2 predict a premature stop codon in all three alleles detected in EIF2AK1 KO cells. (**E**) HeLa (wild-type, EIF2AK1 KO Clone A2 and A3 cell extracts were resolved by SDS–PAGE and subjected to Western blotting with EIF2AK1 polyclonal antibody. (**F**) Wild-type (WT) HeLa cells and two independent EIF2AK1 knockout HeLa cells clones A2, A3 that had all been through in parallel,single-cell sorting, and expansion were treated with OA for 12 hr. Cells were lysed and immunoblotted for Ub pS65, PINK1, EIF2AK1, OPA1, GAPDH and the immunoblots were developed using the LI-COR Odyssey CLx Western Blot imaging system.

**Fig. S10. Analysis of EIF2AK1 expression in S-HeLa PINK1 knockout cells.**
Wild type (WT) and PINK1 knockout S-HeLa cells were treated with OA for 6 and 12 hr. Immunoblot of EIF2AK1, EIF2AK2, EIF2AK3, EIF2AK4, Ub pS65, PINK1, Rab8A, OPA1 and GAPDH were developed using the LI-COR Odyssey CLx Western Blot imaging system.

**Fig. S11. Transcriptional regulation of PINK1 and ATF4 in EIF2AK1 silenced cells.**
(**A**) Relative mRNA levels of PINK1 measured by RT-PCR in HeLa cells treated with OA for 1, 3, 6, 9 and 12 hr. (**B**) as in A, Relative mRNA levels of ATF4 measured by RT-PCR in HeLa cells treated with OA for 1, 3, 6, 9 and 12 hr. (**C**) Relative mRNA levels of ATF4 measured by RT-PCR in Hela cells treated with non-targeting control siRNA pool (NT), siRNA pool targeting PINK1 or, EIF2AK1for 72 hr and with OA in the last 12 hr. (**D**) Relative mRNA levels of EIF2AK1 measured by RT-PCR in Hela cells treated with non-targeting control siRNA pool (NT), siRNA pool targeting PINK1 or, EIF2AK1for 72 hr and with OA in the last 12 hr. (**E**) Relative mRNA levels of EIF2AK2 measured by RT-PCR in HeLa cells treated with non-targeting control siRNA pool (NT), siRNA pool targeting PINK1 or, EIF2AK1for 72

hr and with OA in the last 12 hr. (**F**) Immunoblot analysis for HeLa cells treated with non-targeting control siRNA pool (NT), siRNA pool targeting PINK1 or, EIF2AK1for 72 hr. Co-treated with OA and translation inhibitors DRB and alpha-Amanitib or, OA alone in the last 12 hr. Immunoblot of Ub pS65, PINK1, EIF2AK1, OPA1, GAPDH, ATF4, RNA POLII phospho-S5 (DRB treatment control) and total protein as visualized by PonceauS staining.
Data information: (**A-E**) All data are mean ± SEM; Statistical significance is displayed as $*P \leq 0.05$; $**P \leq 0.01$; $***P \leq 0.001$; $****P \leq 0.0001$; ns, not significant. n = 3 biological replicates. (**A**) one-way ANOVA, Tukey's multiple comparisons test, (**B**) one-way ANOVA, Sidak's multiple comparisons test and (**C-E**) two-way ANOVA, Uncorrected Fisher's LSD multiple comparisons test.

**Fig. S12. Quantitation of translation inhibition of PINK1 upon CHX treatment**
(**A**, **B**) Quantification for Ub pS65/GAPDH and PINK1/GAPDH ratio normalised to the ratio in OA alone samples for Fig. 4E using the Licor Image Studio software.
Data information: (A and B) All data are mean ± SEM; Statistical significance is displayed as $*P \leq 0.05$; $**P \leq 0.01$; $***P \leq 0.001$; $****P \leq 0.0001$; ns, not significant. n = 3 biological replicates, one-way ANOVA, Tukey's multiple comparisons test.

**Fig. S13. Validation of chemical inhibition of the ISR by ISRIB in HeLa and ARPE-19 cells.**
(**A**) HeLa cells seeded in a 10 cm dish, were treated with 300nmol of ISRIB for 24 hr with and without co-treatment of OA for 12 hr or, OA alone for 12 hr. Representative Immunoblot of Ub pS65, PINK1, EIF2AK1, ATF4, OPA1, GAPDH and total protein as visualized by PonceauS staining. (**B**) As in A, ARPE-19 cells were co-treated with 300nmol of ISRIB for 24 hr with OA for 12 hr or, OA alone for 12 hr. Representative Immunoblot of Ub pS65, PINK1, EIF2AK1, ATF4, OPA1, GAPDH and total protein as visualized by PonceauS staining.

**Fig. S14. Investigation of role of EIF2AK1 in DFP-induced mitophagy in cells**
(**A**,**B**) Representative immunoblots (**A**) and quantifications (**B**) of the indicated proteins in lysates of ARPE-19 cells stably expressing the *mito*-QC reporter and HA-Parkin, transfected with non-targeting siRNA (NT) or siRNA targeting EIF2AK1. 2 days post transfection, cells were treated with 1 mM DFP and/or with 50 nM Bafilomycin A (BafA) for an additional 24 hr.
(**C**,**D**) Flow cytometry analysis of mitophagy in cells treated as in A.(**C**)Representative dot plots are shown after analysing GFP and mCherry signals. The percentage of cell underdoing mitophagy is indicated in bold red on each dot plot.
Data information: Overall data are mean ± s.d.; Statistical significance is displayed as $*P \leq 0.05$; $**P \leq 0.01$; $***P \leq 0.001$; $****P \leq 0.0001$; ns, not significant. (**B**) n = 3 biological replicates, two-way ANOVA, Tukey's multiple comparisons test. (**D**) n = 3 biological replicates, two-way ANOVA, Sidak's multiple comparisons test.

### Figure S1

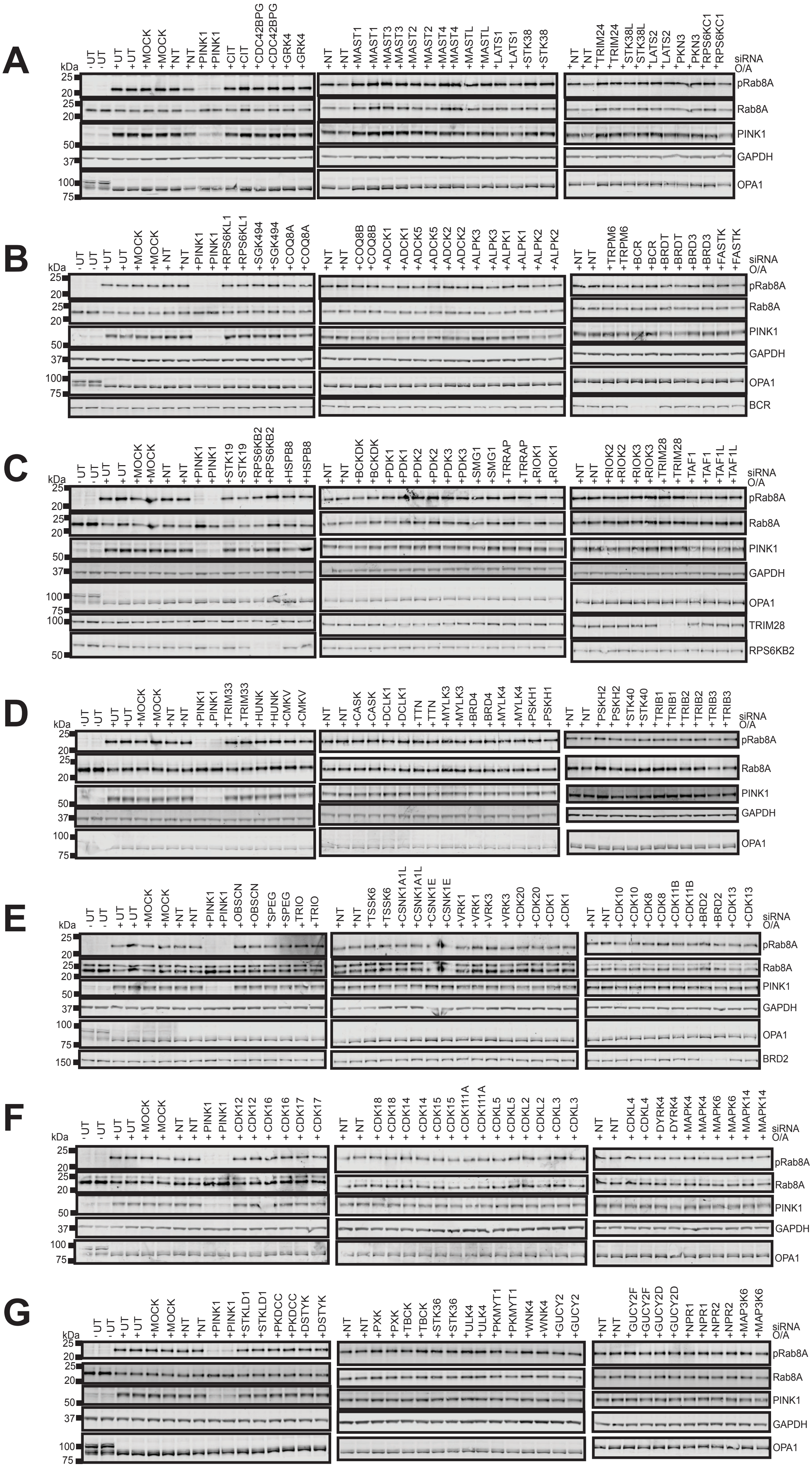

### Figure S2

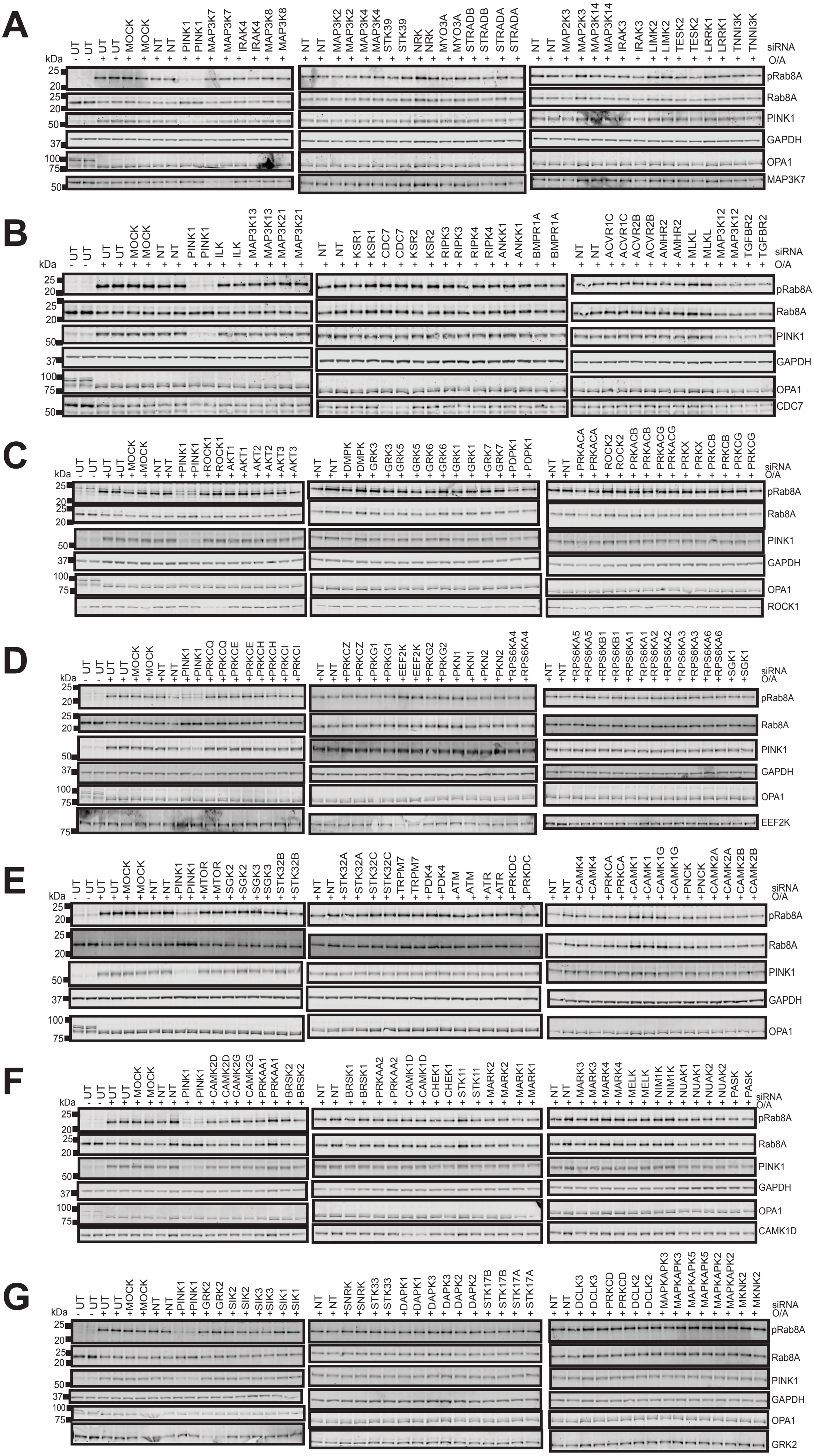
A
- UT, - UT, + UT, + UT, + MOCK, + MOCK, + NT, + NT, + PINK1, + PINK1, + MAP3K7, + MAP3K7, + IRAK4, + IRAK4, + MAP3K8, + MAP3K8
+ NT, + NT, + MAP3K2, + MAP3K2, + MAP3K4, + MAP3K4, + STK39, + STK39, + NRK, + NRK, + MYO3A, + MYO3A, + STRADB, + STRADB, + STRADA, + STRADA
+ NT, + NT, + MAP2K3, + MAP2K3, + MAP3K14, + MAP3K14, + IRAK3, + IRAK3, + LIMK2, + LIMK2, + TESK2, + TESK2, + LRRK1, + LRRK1, + TNNI3K, + TNNI3K
siRNA
O/A
kDa
25
20
25
20
50
37
100
75
50
pRab8A
Rab8A
PINK1
GAPDH
OPA1
MAP3K7
B
- UT, - UT, + UT, + UT, + MOCK, + MOCK, + NT, + NT, + PINK1, + PINK1, + ILK, + ILK, + MAP3K13, + MAP3K13, + MAP3K21, + MAP3K21
+ NT, + NT, + KSR1, + KSR1, + CDC7, + CDC7, + KSR2, + KSR2, + RIPK3, + RIPK3, + RIPK4, + RIPK4, + ANKK1, + ANKK1, + BMPR1A, + BMPR1A
+ NT, + NT, + ACVR1C, + ACVR1C, + ACVR2B, + ACVR2B, + AMHR2, + AMHR2, + MLKL, + MLKL, + MAP3K12, + MAP3K12, + TGFBR2, + TGFBR2
siRNA
O/A
kDa
25
20
25
20
50
37
100
75
50
pRab8A
Rab8A
PINK1
GAPDH
OPA1
CDC7
C
- UT, - UT, + UT, + UT, + MOCK, + MOCK, + NT, + NT, + PINK1, + PINK1, + ROCK1, + ROCK1, + AKT1, + AKT1, + AKT2, + AKT2, + AKT3, + AKT3
+ NT, + NT, + DMPK, + DMPK, + GRK3, + GRK3, + GRK5, + GRK5, + GRK6, + GRK6, + GRK1, + GRK1, + GRK7, + GRK7, + PDPK1, + PDPK1
+ NT, + NT, + PRKACA, + PRKACA, + ROCK2, + ROCK2, + PRKACB, + PRKACB, + PRKACG, + PRKACG, + PRKX, + PRKX, + PRKCB, + PRKCB, + PRKCG, + PRKCG
siRNA
O/A
kDa
25
20
25
20
50
37
100
75
pRab8A
Rab8A
PINK1
GAPDH
OPA1
ROCK1
D
- UT, - UT, + UT, + UT, + MOCK, + MOCK, + NT, + NT, + PINK1, + PINK1, + PRKCQ, + PRKCQ, + PRKCE, + PRKCE, + PRKCH, + PRKCH, + PRKCI, + PRKCI
+ NT, + NT, + PRKCZ, + PRKCZ, + PRKG1, + PRKG1, + EEF2K, + EEF2K, + PRKG2, + PRKG2, + PKN1, + PKN1, + PKN2, + PKN2, + RPS6KA4, + RPS6KA4
+ NT, + NT, + RPS6KA5, + RPS6KA5, + RPS6KB1, + RPS6KB1, + RPS6KA1, + RPS6KA2, + RPS6KA2, + RPS6KA3, + RPS6KA3, + RPS6KA6, + RPS6KA6, + SGK1, + SGK1
siRNA
O/A
kDa
25
20
25
20
50
37
100
75
75
pRab8A
Rab8A
PINK1
GAPDH
OPA1
EEF2K
E
- UT, - UT, + UT, + UT, + MOCK, + MOCK, + NT, + NT, + PINK1, + PINK1, + MTOR, + MTOR, + SGK2, + SGK2, + SGK3, + SGK3, + STK32B, + STK32B
+ NT, + NT, + STK32A, + STK32A, + STK32C, + STK32C, + TRPM7, + TRPM7, + PDK4, + PDK4, + ATM, + ATM, + ATR, + ATR, + PRKDC, + PRKDC
+ NT, + NT, + CAMK4, + CAMK4, + PRKCA, + PRKCA, + CAMK1, + CAMK1, + CAMK1G, + CAMK1G, + PNCK, + PNCK, + CAMK2A, + CAMK2A, + CAMK2B, + CAMK2B
siRNA
O/A
kDa
25
20
25
20
50
37
100
75
pRab8A
Rab8A
PINK1
GAPDH
OPA1
F
- UT, - UT, + UT, + UT, + MOCK, + MOCK, + NT, + NT, + PINK1, + PINK1, + CAMK2D, + CAMK2D, + CAMK2G, + CAMK2G, + PRKAA1, + PRKAA1, + BRSK2, + BRSK2
+ NT, + NT, + BRSK1, + BRSK1, + PRKAA2, + PRKAA2, + CAMK1D, + CAMK1D, + CHEK1, + CHEK1, + STK11, + STK11, + MARK2, + MARK2, + MARK1, + MARK1
+ NT, + NT, + MARK3, + MARK3, + MARK4, + MARK4, + MELK, + MELK, + NIM1K, + NIM1K, + NUAK1, + NUAK1, + NUAK2, + NUAK2, + PASK, + PASK
siRNA
O/A
kDa
25
20
25
20
50
37
100
75
pRab8A
Rab8A
PINK1
GAPDH
OPA1
CAMK1D
G
- UT, - UT, + UT, + UT, + MOCK, + MOCK, + NT, + NT, + PINK1, + PINK1, + GRK2, + GRK2, + SIK2, + SIK2, + SIK3, + SIK3, + SIK1, + SIK1
+ NT, + NT, + SNRK, + SNRK, + STK33, + STK33, + DAPK1, + DAPK1, + DAPK3, + DAPK3, + DAPK2, + DAPK2, + STK17B, + STK17B, + STK17A, + STK17A
+ NT, + NT, + DCLK3, + DCLK3, + PRKCD, + PRKCD, + DCLK2, + DCLK2, + MAPKAPK3, + MAPKAPK3, + MAPKAPK5, + MAPKAPK5, + MAPKAPK2, + MAPKAPK2, + MKNK2, + MKNK2
siRNA
O/A
kDa
25
20
25
20
50
37
100
75
pRab8A
Rab8A
PINK1
GAPDH
OPA1
GRK2

### Figure S3

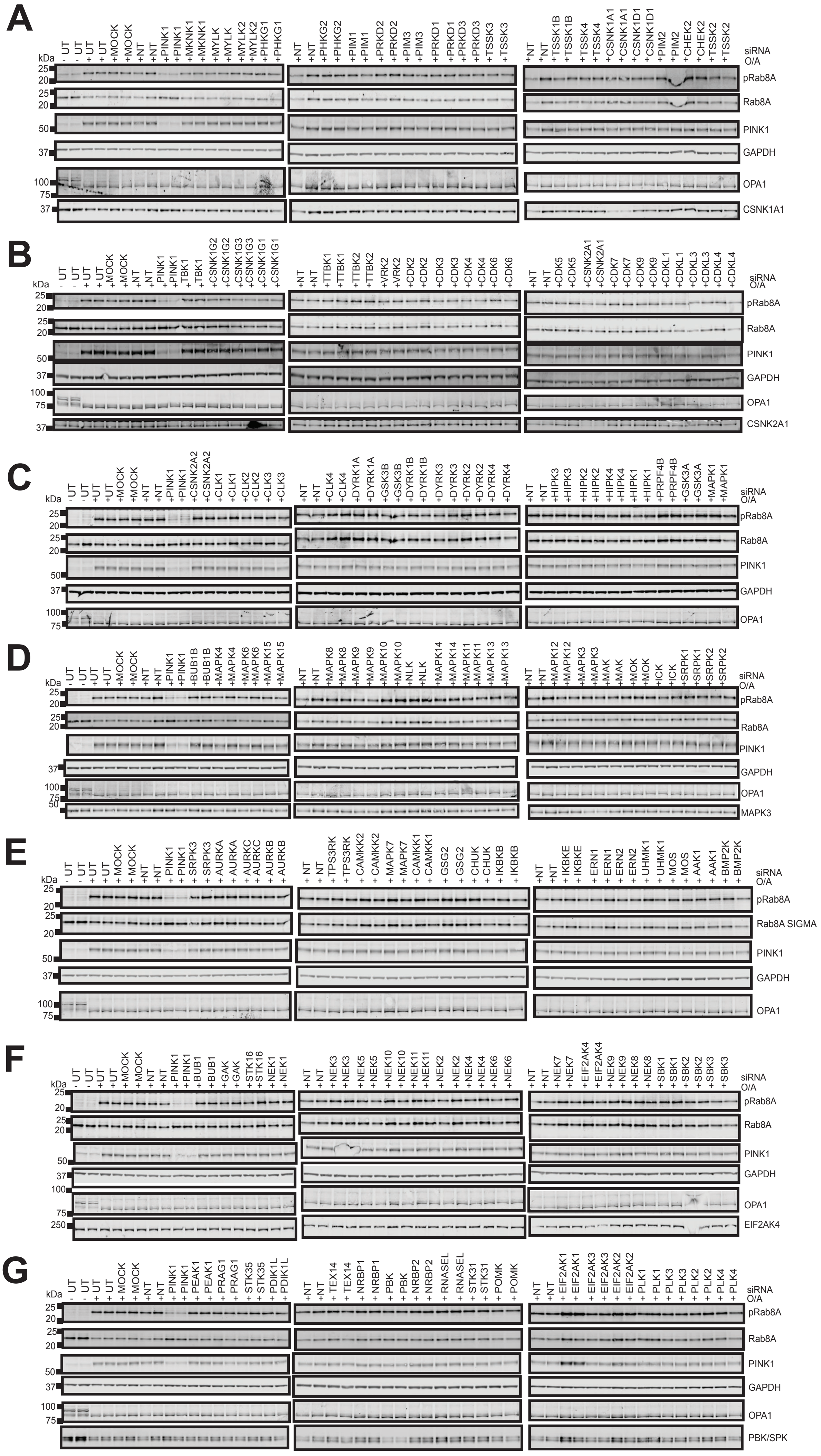
A
kDa
- UT - UT + UT +MOCK +MOCK +NT +NT +PINK1 +PINK1 +MKNK1 +MKNK1 +MYLK +MYLK +MYLK2 +MYLK2 +PHKG1 +PHKG1
+NT +NT +PHKG2 +PHKG2 +PIM1 +PIM1 +PRKD2 +PRKD2 +PIM3 +PIM3 +PRKD1 +PRKD1 +PRKD3 +PRKD3 +TSSK3 +TSSK3
+NT +NT +TSSK1B +TSSK1B +TSSK4 +TSSK4 +CSNK1A1 +CSNK1A1 +CSNK1D1 +CSNK1D1 +PIM2 +PIM2 +CHEK2 +CHEK2 +TSSK2 +TSSK2
siRNA
O/A
pRab8A
Rab8A
PINK1
GAPDH
OPA1
CSNK1A1
B
- UT - UT + UT +MOCK +MOCK +NT +NT +PINK1 +PINK1 +TBK1 +TBK1 +CSNK1G2 +CSNK1G2 +CSNK1G3 +CSNK1G3 +CSNK1G1 +CSNK1G1
+NT +NT +TTBK1 +TTBK1 +TTBK2 +TTBK2 +VRK2 +VRK2 +CDK2 +CDK2 +CDK3 +CDK3 +CDK4 +CDK4 +CDK6 +CDK6
+NT +NT +CDK5 +CDK5 +CSNK2A1 +CSNK2A1 +CDK7 +CDK7 +CDK9 +CDK9 +CDKL1 +CDKL1 +CDKL3 +CDKL3 +CDKL4 +CDKL4
pRab8A
Rab8A
PINK1
GAPDH
OPA1
CSNK2A1
C
- UT - UT + UT +MOCK +MOCK +NT +NT +PINK1 +PINK1 +CSNK2A2 +CSNK2A2 +CLK1 +CLK1 +CLK2 +CLK2 +CLK3 +CLK3
+NT +NT +CLK4 +CLK4 +DYRK1A +DYRK1A +GSK3B +GSK3B +DYRK1B +DYRK1B +DYRK3 +DYRK3 +DYRK2 +DYRK2 +DYRK4 +DYRK4
+NT +NT +HIPK3 +HIPK3 +HIPK2 +HIPK2 +HIPK4 +HIPK4 +HIPK1 +HIPK1 +PRPF4B +PRPF4B +GSK3A +GSK3A +MAPK1 +MAPK1
pRab8A
Rab8A
PINK1
GAPDH
OPA1
D
- UT - UT + UT +MOCK +MOCK +NT +NT +PINK1 +PINK1 +BUB1B +BUB1B +MAPK4 +MAPK4 +MAPK6 +MAPK6 +MAPK15 +MAPK15
+NT +NT +MAPK8 +MAPK8 +MAPK9 +MAPK9 +MAPK10 +MAPK10 +NLK +NLK +MAPK14 +MAPK14 +MAPK11 +MAPK11 +MAPK13 +MAPK13
+NT +NT +MAPK12 +MAPK12 +MAPK3 +MAPK3 +MAK +MAK +MOK +MOK +ICK +ICK +SRPK1 +SRPK1 +SRPK2 +SRPK2
pRab8A
Rab8A
PINK1
GAPDH
OPA1
MAPK3
E
- UT - UT + UT + UT +MOCK +MOCK +NT +NT + PINK1 + PINK1 + SRPK3 + SRPK3 + AURKA + AURKA + AURKC + AURKC + AURKB + AURKB
+ NT + NT + TPS3RK + TPS3RK + CAMKK2 + CAMKK2 + MAPK7 + MAPK7 + CAMKK1 + CAMKK1 + GSG2 + GSG2 + CHUK + CHUK + IKBKB + IKBKB
+NT +NT + IKBKE + IKBKE + ERN1 + ERN1 + ERN2 + ERN2 + UHMK1 + UHMK1 + MOS + MOS + AAK1 + AAK1 + BMP2K + BMP2K
pRab8A
Rab8A SIGMA
PINK1
GAPDH
OPA1
F
- UT - UT + UT +MOCK +MOCK +NT +NT +PINK1 +PINK1 +BUB1 +BUB1 +GAK +GAK +STK16 +STK16 +NEK1 +NEK1
+NT +NT +NEK3 +NEK3 +NEK5 +NEK5 +NEK10 +NEK10 +NEK11 +NEK11 +NEK2 +NEK2 +NEK4 +NEK4 +NEK6 +NEK6
+NT +NT +NEK7 +NEK7 + EIF2AK4 + EIF2AK4 +NEK9 +NEK9 +NEK8 +NEK8 +SBK1 +SBK1 +SBK2 +SBK2 +SBK3 +SBK3
pRab8A
Rab8A
PINK1
GAPDH
OPA1
EIF2AK4
G
- UT - UT + UT + MOCK + MOCK + NT + NT + PINK1 + PINK1 + PEAK1 + PEAK1 + PRAG1 + PRAG1 + STK35 + STK35 + PDIK1L + PDIK1L
+NT +NT + TEX14 + TEX14 + NRBP1 + NRBP1 + PBK + PBK + NRBP2 + NRBP2 + RNASEL + RNASEL + STK31 + STK31 + POMK + POMK
+NT +NT + EIF2AK1 + EIF2AK1 + EIF2AK3 + EIF2AK3 + EIF2AK2 + EIF2AK2 + PLK1 + PLK1 + PLK3 + PLK3 + PLK2 + PLK2 + PLK4 + PLK4
pRab8A
Rab8A
PINK1
GAPDH
OPA1
PBK/SPK

### Figure S4

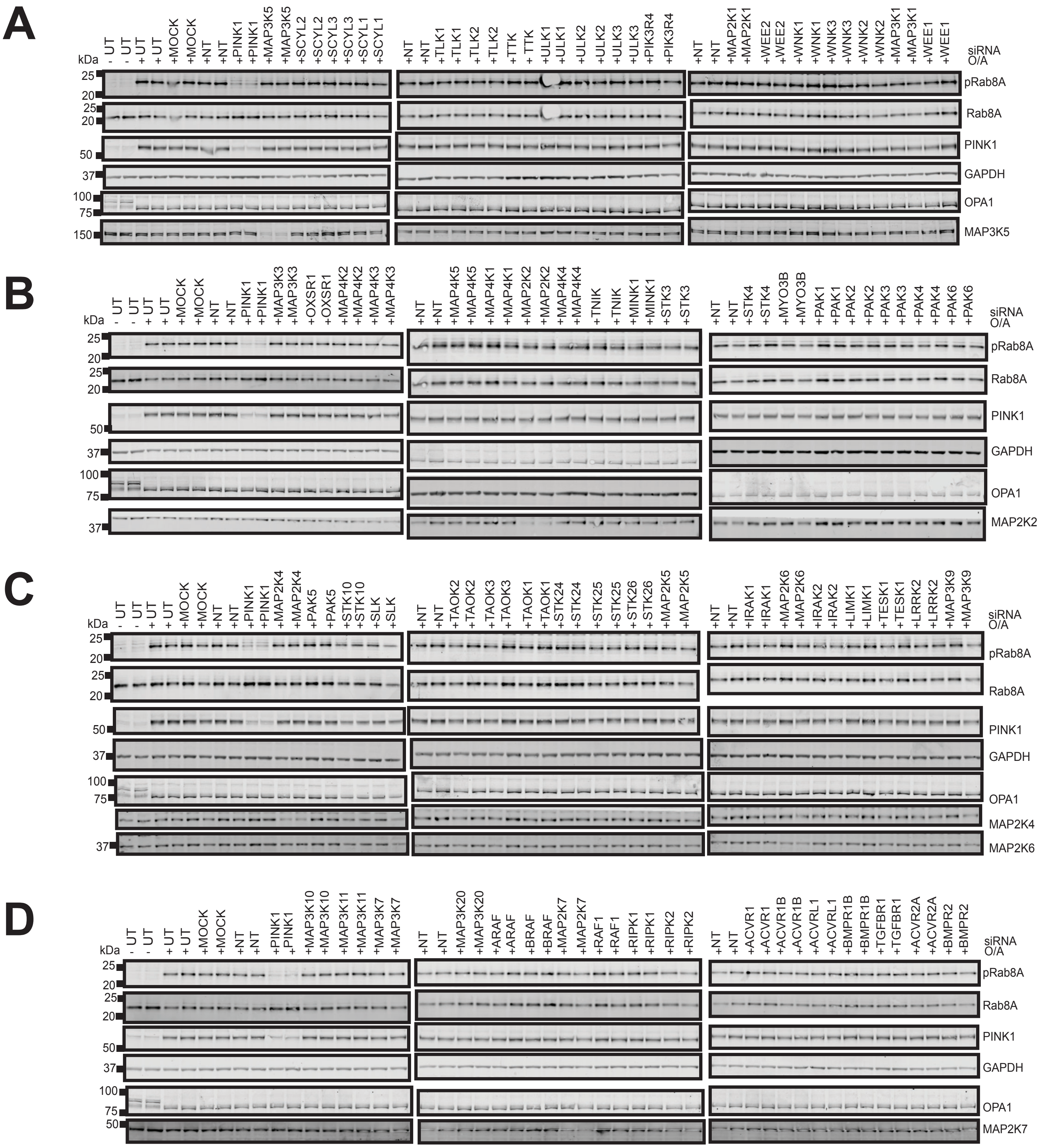
A
B
C
D
kDa
siRNA
O/A
pRab8A
Rab8A
PINK1
GAPDH
OPA1
MAP3K5
MAP2K2
MAP2K4
MAP2K6
MAP2K7

### Figure S5

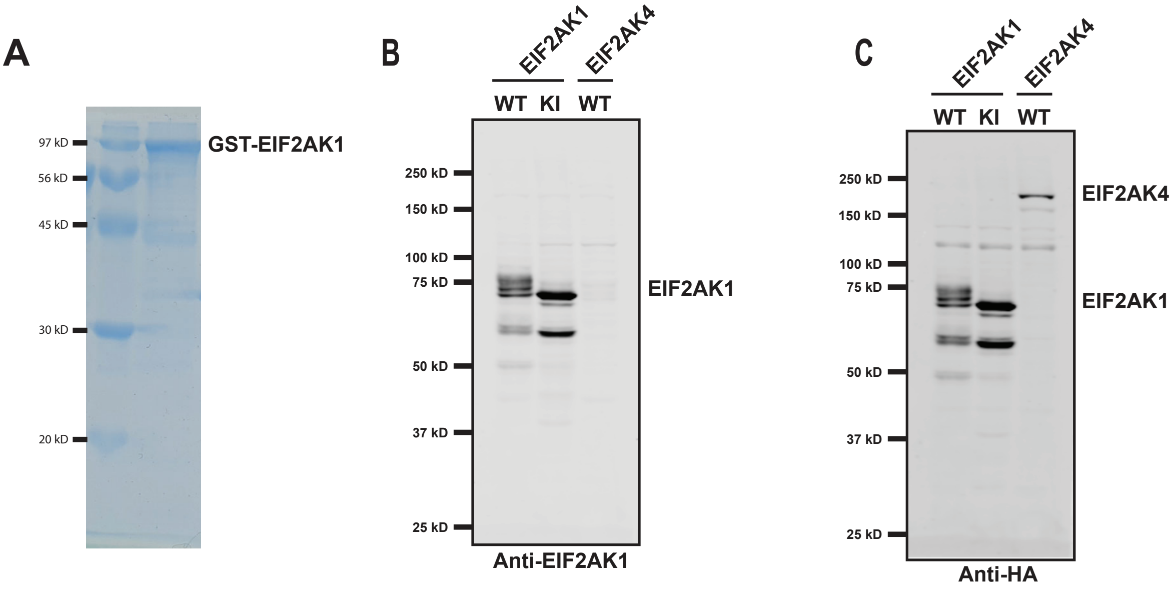
A
97 kD
56 kD
45 kD
30 kD
20 kD
GST-EIF2AK1
B
EIF2AK1
EIF2AK4
WT
KI
WT
250 kD
150 kD
100 kD
75 kD
50 kD
37 kD
25 kD
EIF2AK1
Anti-EIF2AK1
C
EIF2AK1
EIF2AK4
WT
KI
WT
250 kD
150 kD
100 kD
75 kD
50 kD
37 kD
25 kD
EIF2AK4
EIF2AK1
Anti-HA

### Figure S6

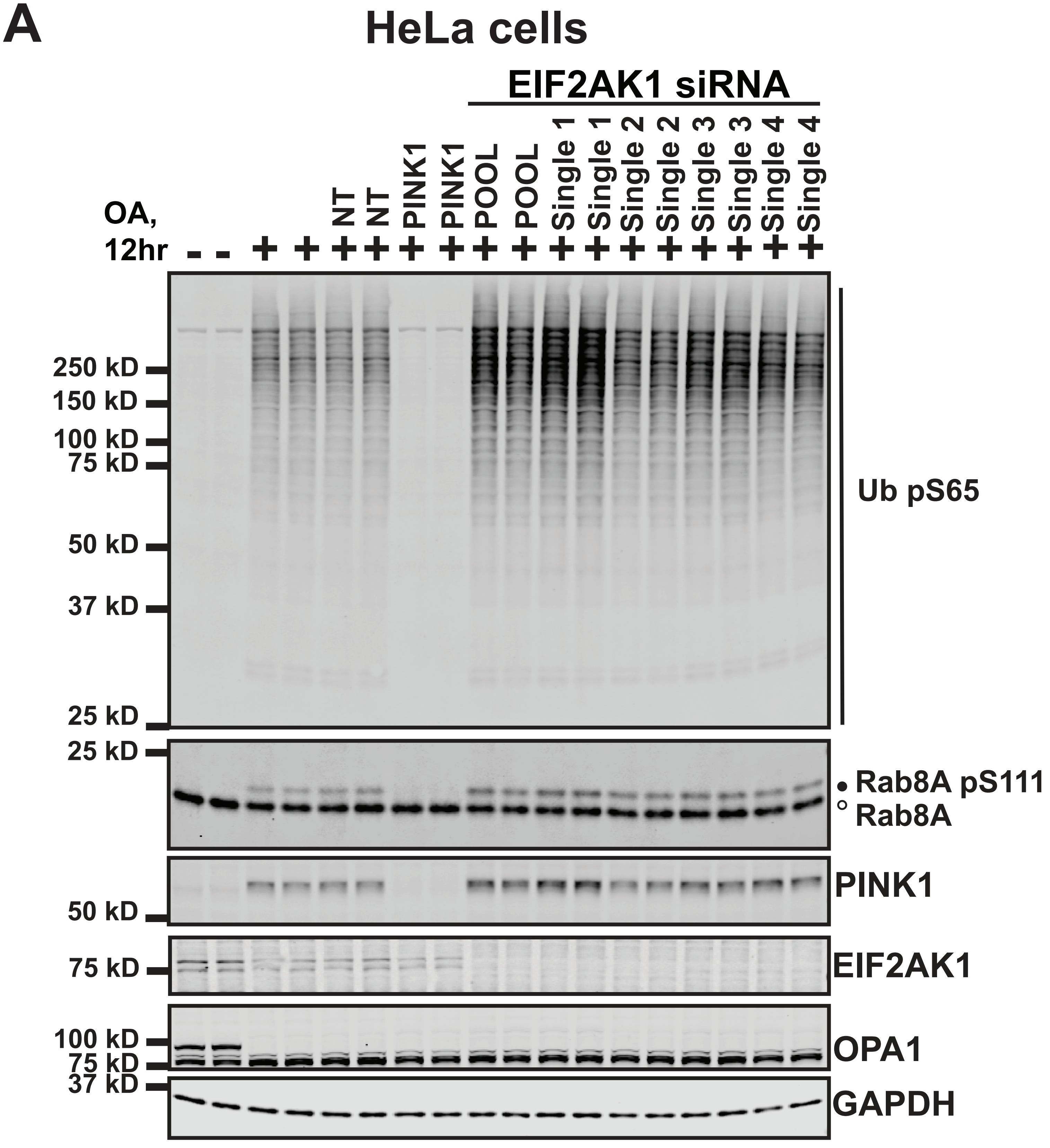
A
HeLa cells
EIF2AK1 siRNA
OA, 12hr
- - + + + + + + + + + + + + + + + +
NT
NT
PINK1
PINK1
POOL
POOL
Single 1
Single 1
Single 2
Single 2
Single 3
Single 3
Single 4
Single 4
250 kD
150 kD
100 kD
75 kD
50 kD
37 kD
25 kD
Ub pS65
25 kD
Rab8A pS111
Rab8A
PINK1
50 kD
75 kD
EIF2AK1
100 kD
75 kD
OPA1
37 kD
GAPDH

### Figure S7

A

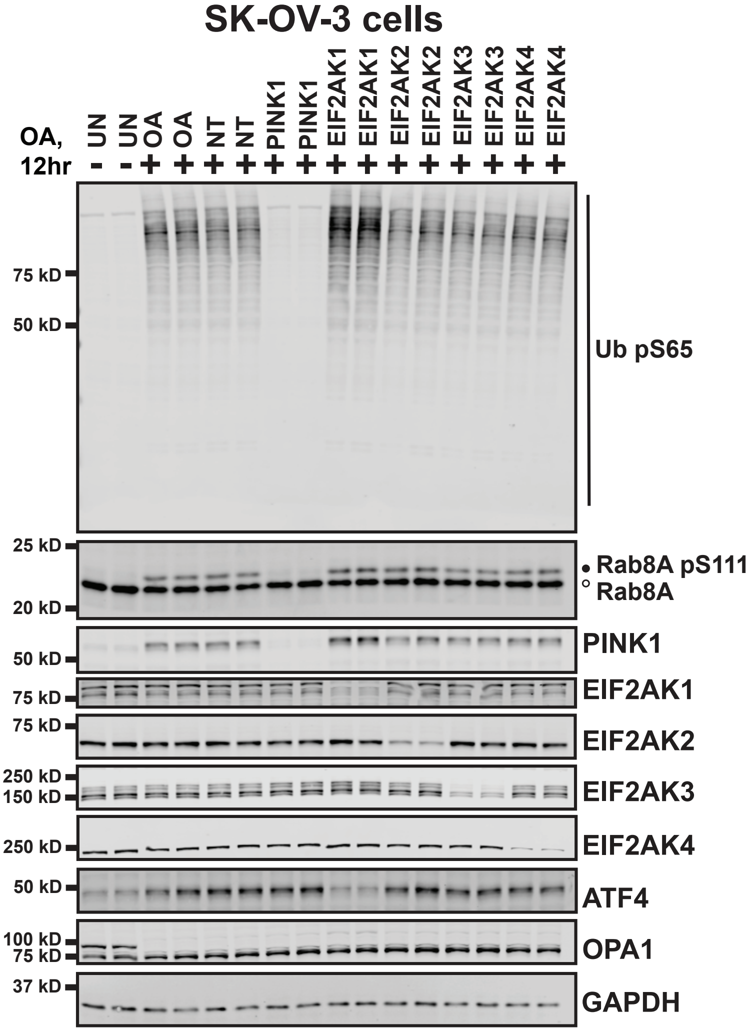


B

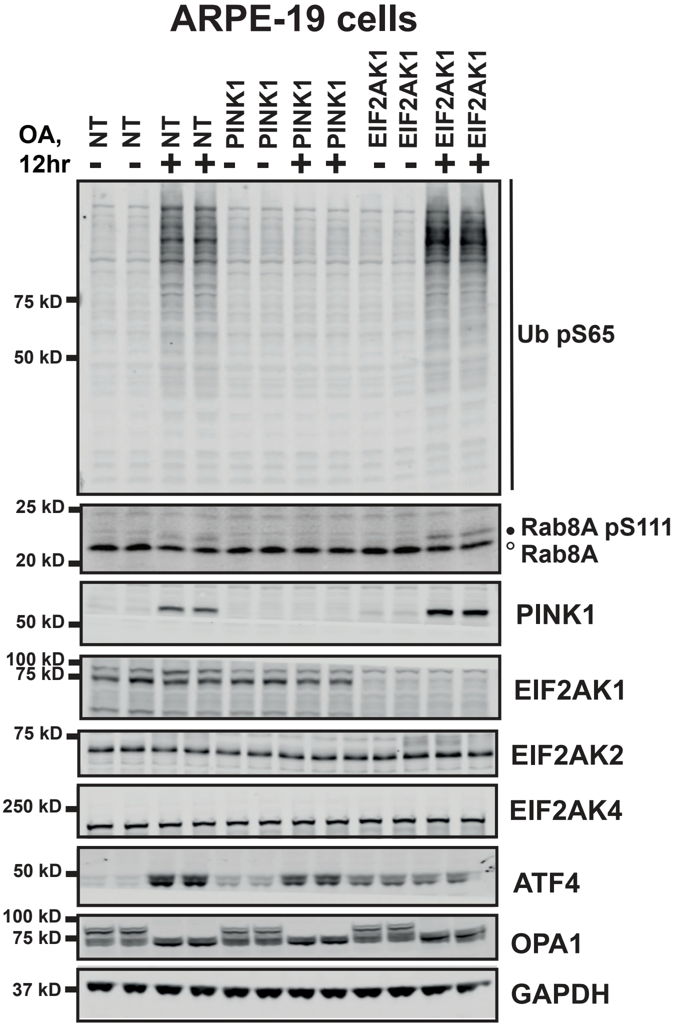


C

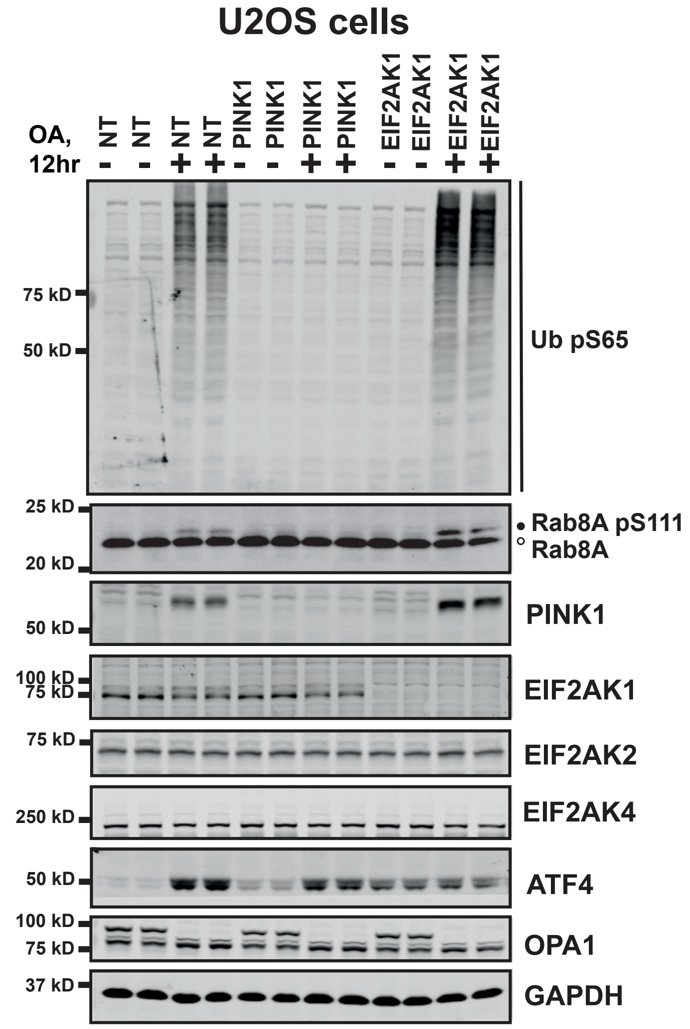

### Figure S8

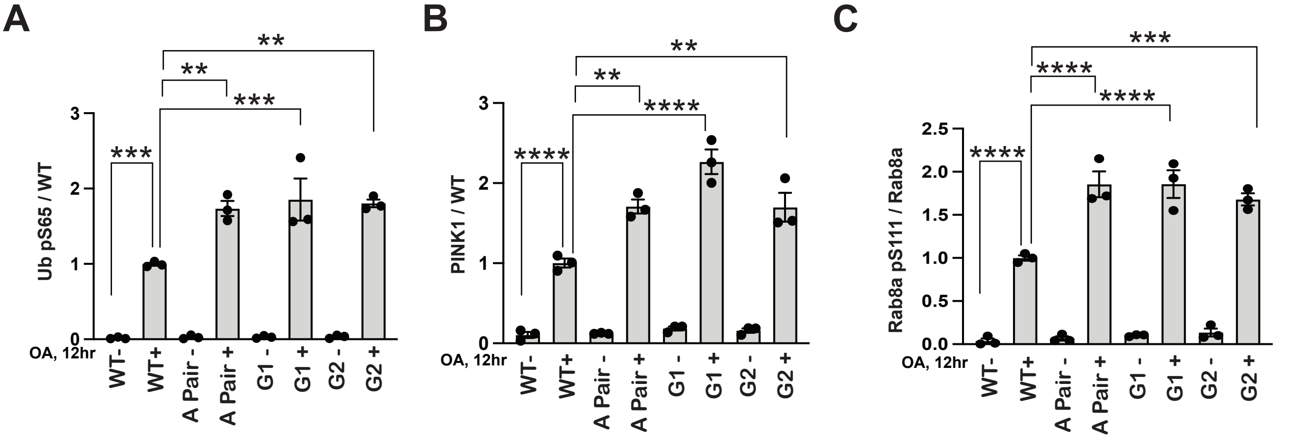
A
Ub pS65 / WT
0
1
2
3
OA, 12hr
WT-
WT+
A Pair -
A Pair +
G1 -
G1 +
G2 -
G2 +
***
**
***
**
B
PINK1 / WT
0
1
2
3
OA, 12hr
WT-
WT+
A Pair -
A Pair +
G1 -
G1 +
G2 -
G2 +
****
**
****
**
C
Rab8a pS111 / Rab8a
0.0
0.5
1.0
1.5
2.0
2.5
OA, 12hr
WT-
WT+
A Pair -
A Pair +
G1 -
G1 +
G2 -
G2 +
****
****
****
***

#### Figure S9

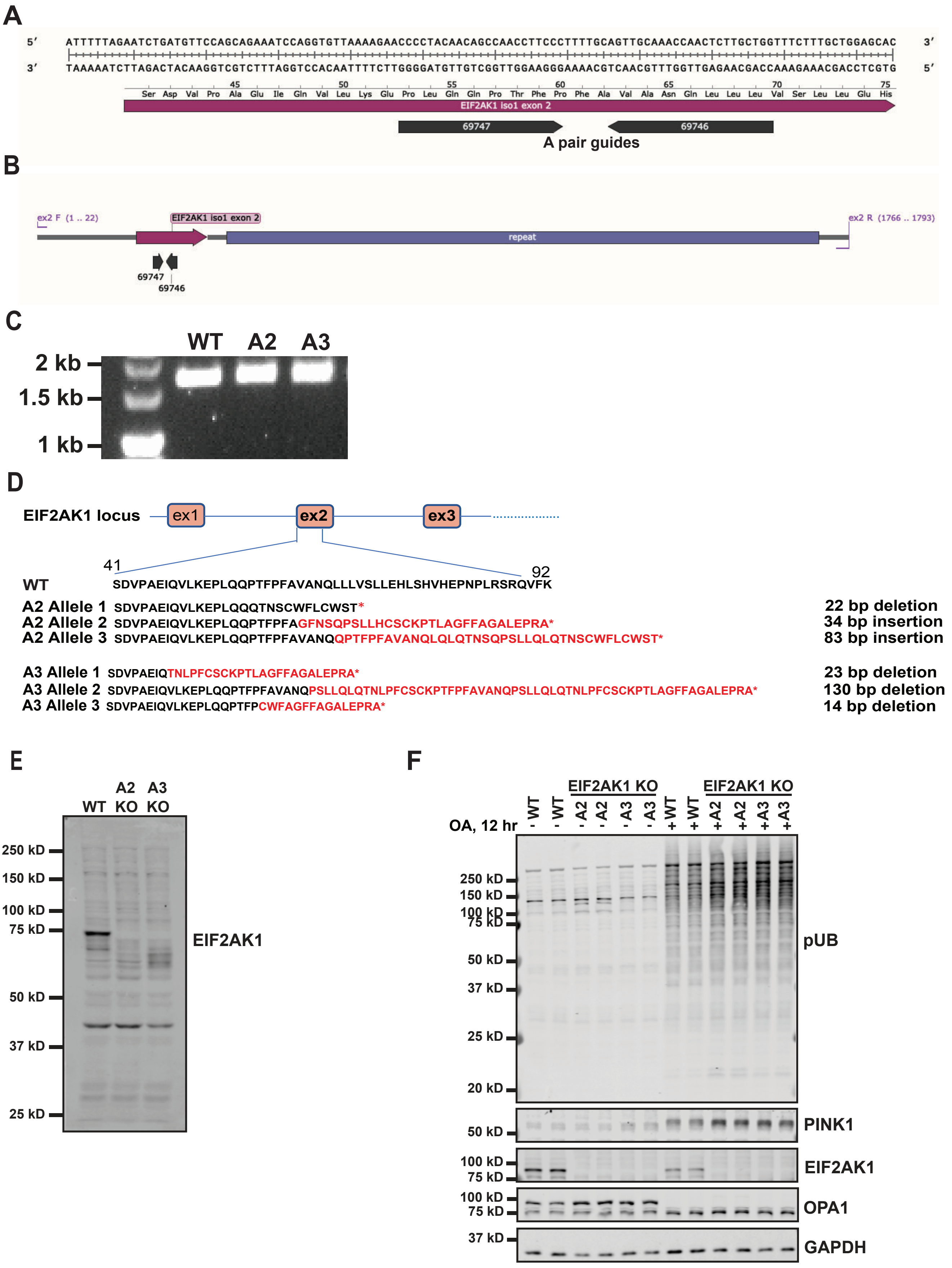

A
5' 3'
3' 5'
Ser Asp Val Pro Ala Glu Ile Gln Val Leu Lys Glu Pro Leu Gln Gln Pro Thr Phe Pro Phe Ala Val Ala Asn Gln Leu Leu Leu Val Ser Leu Glu His
EIF2AK1 iso1 exon 2
69747
69746
A pair guides
B
ex2 F (1 .. 22)
EIF2AK1 iso1 exon 2
repeat
ex2 R (1766 .. 1793)
69747
69746
C
WT A2 A3
2 kb
1.5 kb
1 kb
D
EIF2AK1 locus
ex1
ex2
ex3
41
92
WT SDVPAEIQVLKEPLQQPTFPFAVANQLLLVSLLEHLSHVHEPNPLRSRQVFK
A2 Allele 1 SDVPAEIQVLKEPLQQQTNSCWFLCWST* 22 bp deletion
A2 Allele 2 SDVPAEIQVLKEPLQQPTFPFAGFNSQPSLLHCSCKPTLAGFFAGALEPRA* 34 bp insertion
A2 Allele 3 SDVPAEIQVLKEPLQQPTFPFAVANQQPTFPFAVANQLQLQTNSQPSLLQLQTNSCWFLCWST* 83 bp insertion
A3 Allele 1 SDVPAEIQTNLPFCSCKPTLAGFFAGALEPRA* 23 bp deletion
A3 Allele 2 SDVPAEIQVLKEPLQQPTFPFAVANQPSLLQLQTNLPFCSCKPTFPFAVANQPSLLQLQTNLPFCSCKPTLAGFFAGALEPRA* 130 bp deletion
A3 Allele 3 SDVPAEIQVLKEPLQQPTFPCWFAGFFAGALEPRA* 14 bp deletion
E
WT A2 KO A3 KO
250 kD
150 kD
100 kD
75 kD
50 kD
37 kD
25 kD
EIF2AK1
F
EIF2AK1 KO
EIF2AK1 KO
OA, 12 hr
- WT - WT - A2 - A2 - A3 - A3 + WT + WT + A2 + A2 + A3 + A3
250 kD
150 kD
100 kD
75 kD
50 kD
37 kD
25 kD
20 kD
pUB
50 kD
PINK1
100 kD
75 kD
EIF2AK1
100 kD
75 kD
OPA1
37 kD
GAPDH

### Figure S10

## A

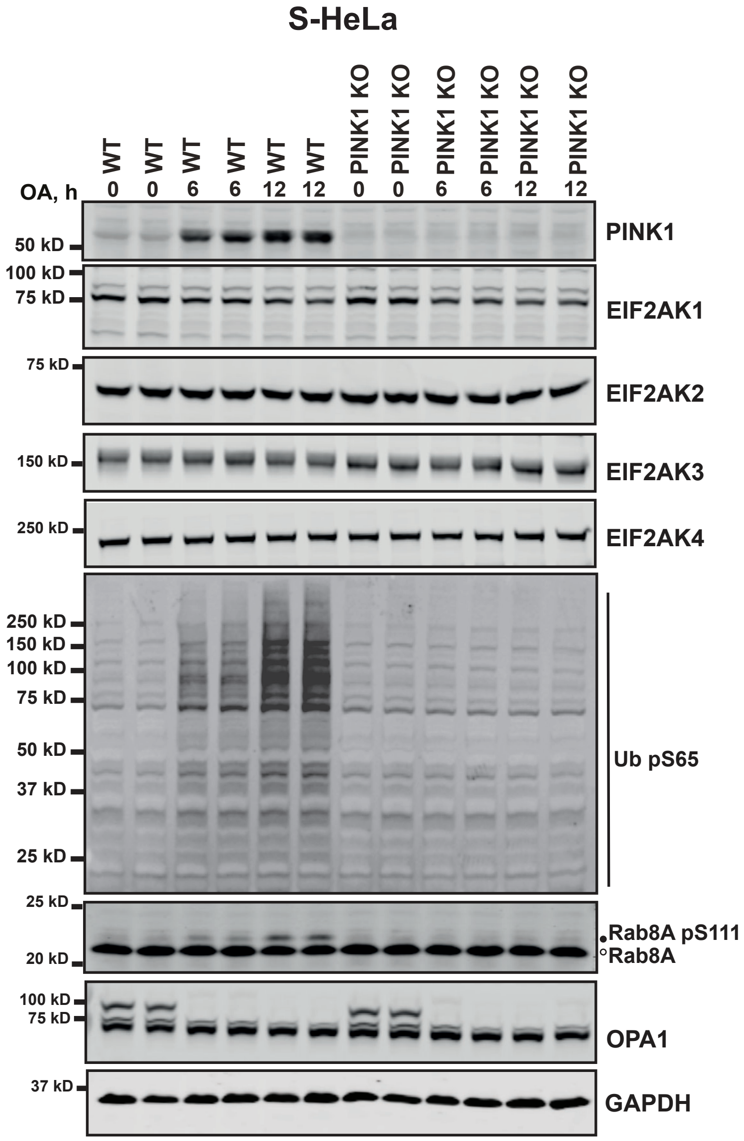
S-HeLa
WT WT WT WT WT WT PINK1 KO PINK1 KO PINK1 KO PINK1 KO PINK1 KO PINK1 KO
OA, h 0 0 6 6 12 12 0 0 6 6 12 12
50 kD
PINK1
100 kD
75 kD
EIF2AK1
75 kD
EIF2AK2
150 kD
EIF2AK3
250 kD
EIF2AK4
250 kD
150 kD
100 kD
75 kD
50 kD
37 kD
25 kD
Ub pS65
25 kD
20 kD
•Rab8A pS111
°Rab8A
100 kD
75 kD
OPA1
37 kD
GAPDH

#### Figure S11

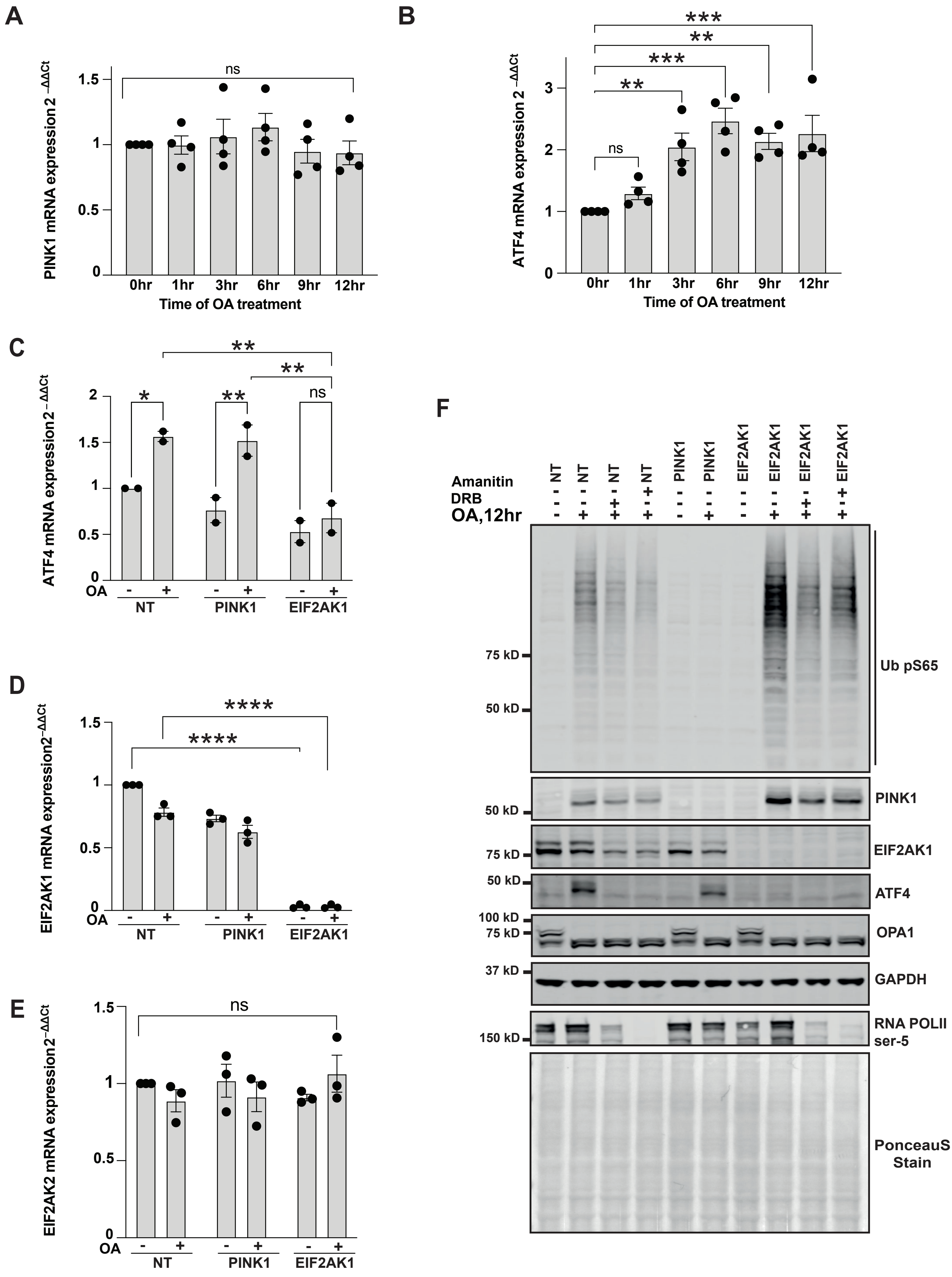
A
PINK1 mRNA expression 2 −ΔΔCt
ns
Time of OA treatment
0hr 1hr 3hr 6hr 9hr 12hr
B
ATF4 mRNA expression 2 −ΔΔCt
ns
**
***
Time of OA treatment
C
ATF4 mRNA expression 2−ΔΔCt
OA
NT
PINK1
EIF2AK1
D
EIF2AK1 mRNA expression2−ΔΔCt
****
E
EIF2AK2 mRNA expression2−ΔΔCt
F
Amanitin
DRB
OA,12hr
NT
PINK1
EIF2AK1
75 kD
50 kD
Ub pS65
PINK1
EIF2AK1
ATF4
100 kD
OPA1
37 kD
GAPDH
150 kD
RNA POLII ser-5
PonceauS Stain

### Figure S12

A

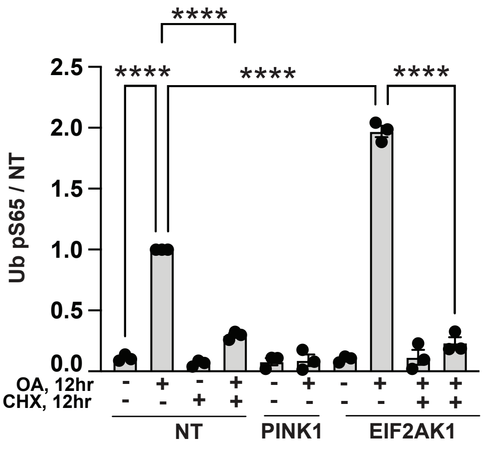


B

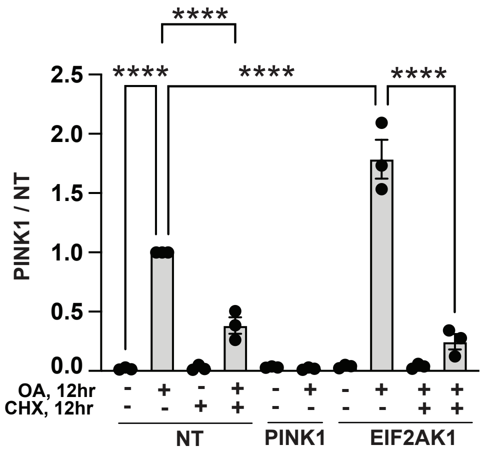

### Figure S13

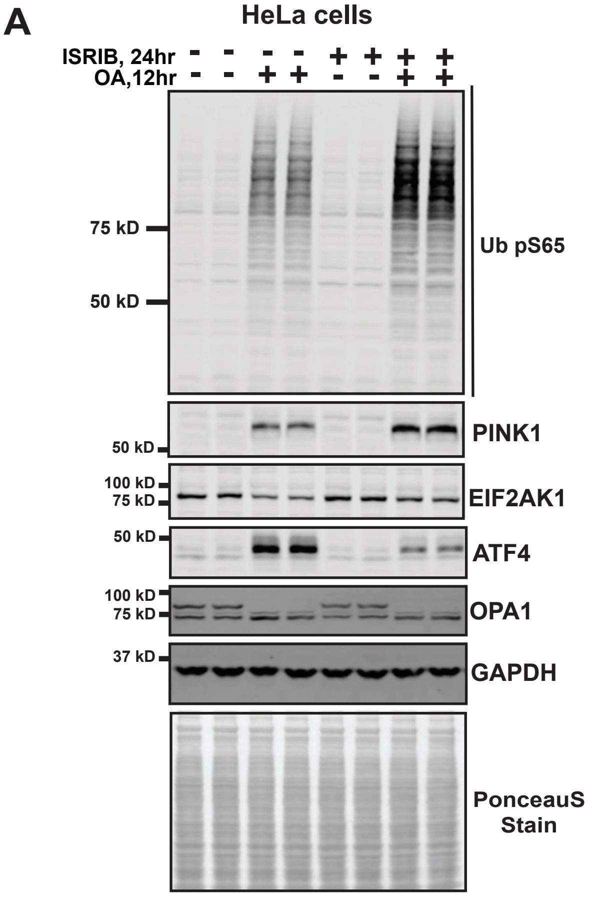
A
HeLa cells
ISRIB, 24hr - - - - + + + +
OA,12hr - - + + - - + +
75 kD
50 kD
Ub pS65
50 kD
PINK1
100 kD
75 kD
EIF2AK1
50 kD
ATF4
100 kD
75 kD
OPA1
37 kD
GAPDH
PonceauS Stain
B
ARPE-19 cells
ISRIB, 24hr - - - - + +
OA,12hr - - + + + +
75 kD
50 kD
Ub pS65
50 kD
PINK1
100 kD
75 kD
EIF2AK1
50 kD
ATF4
100 kD
75 kD
OPA1
37 kD
GAPDH
PonceauS Stain

### Figure S14

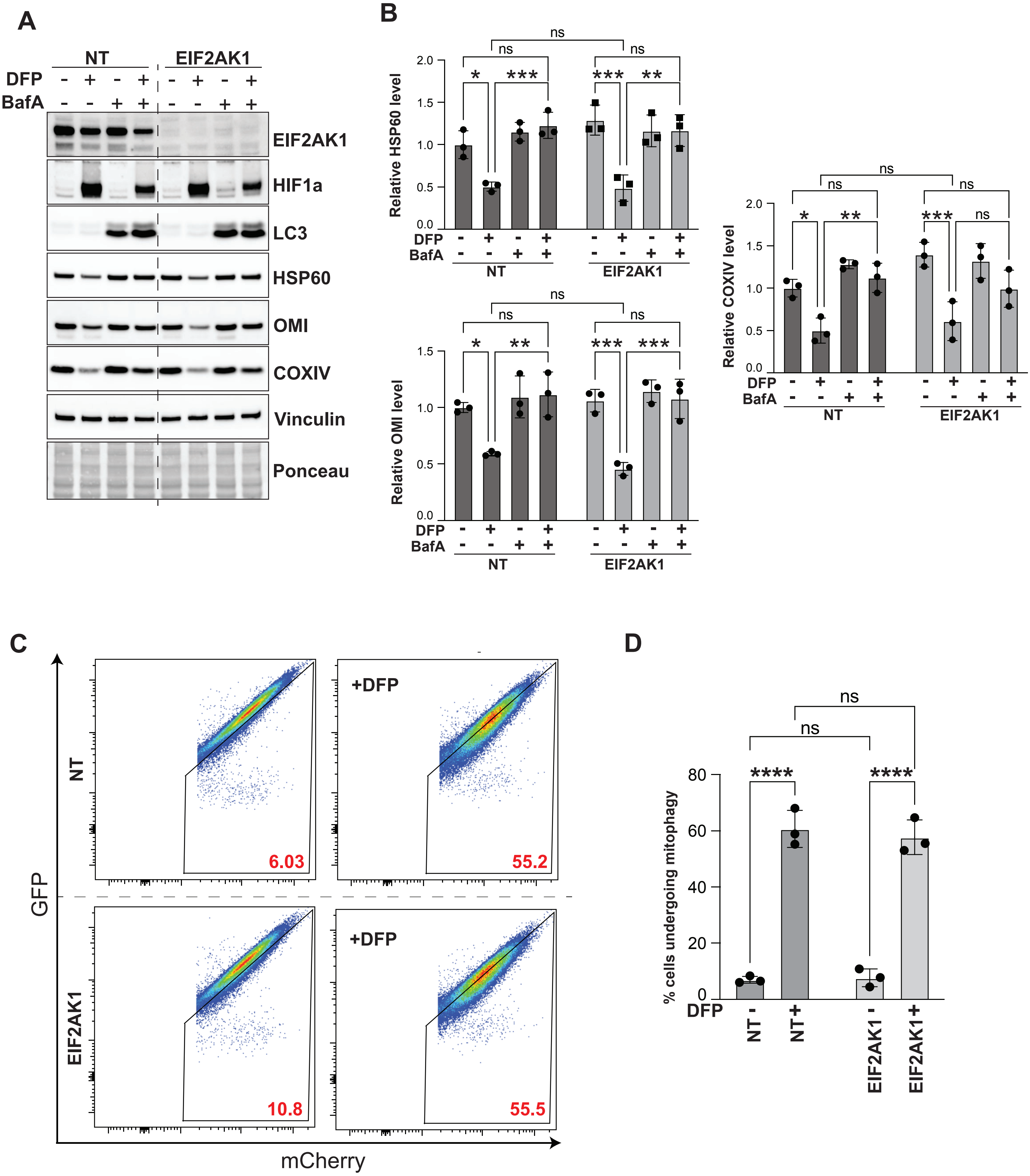
A
NT
EIF2AK1
DFP
BafA
EIF2AK1
HIF1a
LC3
HSP60
OMI
COXIV
Vinculin
Ponceau
B
Relative HSP60 level
Relative OMI level
Relative COXIV level
ns
C
GFP
mCherry
NT
EIF2AK1
+DFP
6.03
55.2
10.8
55.5
D
% cells undergoing mitophagy
NT -
NT +
EIF2AK1 -
EIF2AK1 +
